# Senescence and entrenchment in evolution of amino acid sites

**DOI:** 10.1101/794743

**Authors:** A. V. Stolyarova, E. Nabieva, V. V. Ptushenko, A. V. Favorov, A. V. Popova, A. D. Neverov, G. A. Bazykin

## Abstract

Amino acid propensities at a site change in the course of protein evolution. This may happen for two reasons. Changes may be triggered by substitutions at epistatically interacting sites elsewhere in the genome; alternatively, they may arise due to environmental changes that are external to the genome. Here, we design a framework for distinguishing between these alternatives. Using analytical modelling and simulations, we show that they cause opposite dynamics of the fitness of the allele currently occupying the site: its fitness tends to increase with the time since its origin due to epistasis (“entrenchment”), but to decrease due to random environmental fluctuations (“senescence”). We analyse the phylogenetic distribution of substitutions in nuclear genomes, and show that among the amino acids originating at negatively selected sites of vertebrates, nearly all experience strong entrenchment. By contrast, among the amino acids originating at positively selected sites, 18% experience senescence. A similar pattern is observed in phylogenies of insects. We propose that senescence of the current allele is a cause of adaptive evolution.

## Introduction

Fitness landscape is the key concept of evolutionary biology, and its description is necessary to fully understand adaptive evolution and speciation^1–4^. Unfortunately, the large dimensionality of even the landscapes of individual proteins makes them impossible to measure comprehensively in a direct experiment^5,6^. Still, methods of comparative genomics can be used to assess the integral features of fitness landscapes. The simplest informative unit of landscape structure is the single-position fitness landscape (SPFL)^7^, *i.e.*, a vector of fitness values of all possible alleles at an individual genomic position. SPFLs change with time^8–15^; this may affect the optimality of the allele that is currently prevalent at this site, influencing subsequent evolution.

One factor entailing changes of SPFL is substitutions at other sites of the genome. For this to be the case, these substitutions need to affect the relative fitness of different variants at the considered site, i.e., these sites have to be involved in epistatic interactions. Epistasis has been postulated to be a prevalent factor of protein evolution and divergence across species^2,6,9,16–24^. One expected manifestation of genome-wide epistasis is entrenchment, or the evolutionary Stokes shift^12,25,26^ — a phenomenon whereby the relative fitness of the allele currently prevalent at the site increases as substitutions at interacting sites accumulate. The reason for this increase is the constraint imposed by the site in consideration onto epistatically interacting sites. The evolution of the remaining sequence is constrained to preserve the high fitness of the resident allele, and may even increase it; at the same time, this sequence is free to evolve to become less compatible with other variants not currently present at the site. Over time, this leads to an increase in the fitness of the current allele relative to other alleles, including those that resided at this site earlier. Entrenchment was demonstrated in simulated protein evolution^12,25,26^, and cases when the same allele has originated more than once (homoplasies) follow the phylogenetic distribution indicative of it. In particular, reversals of past substitutions were shown to become more deleterious with time, confirming that the current allele becomes more preferable compared to the previous one^27–30^. The decline in the rate of reversals is caused both by the increase in the fitness of the current allele and the decrease in the fitness of the replaced allele^28^.

However, the SPFL may change due to environmental fluctuations even in the absence of epistasis. If such changes are recurrent, the fitness landscape becomes a time-dependent “seascape”^16,31–36^. This leads to recurrent positive selection (fluctuating selection) in favor of the newly beneficial alleles and to adaptive evolution^35,37–40^. Nowadays, the way fluctuating selection shapes the dynamics of the relative fitness of the current allele remains poorly studied. Here, we characterize the effects of epistasis and of fluctuating selection on SPFL changes and estimate the contribution of these forces in past evolution.

## Results

### Environmental fluctuations decrease the fitness of the current allele

First, we ask how fluctuating selection affects the relative fitness of different alleles at a site. If changes of the SPFL are random with regard to the identity of the allele currently residing at the site, we expect that they, on average, will reduce its relative fitness. Indeed, due to natural selection, the relative fitness conferred by the current variant is, on average, higher than that of a random variant at this site, so that a random change to the SPFL will, on average, reduce it (see formal proof in Supplementary Text 1). An episode of positive selection triggered by an SPFL change may then cause the spread of a novel variant which would confer high fitness till the next SPFL change.

To illustrate this, we simulate amino acid evolution on a randomly changing fitness landscape. In this simulation, the fitness values for each of the 20 possible amino acids are drawn from a predefined distribution, and the amino acid substitutions occur with probabilities determined by the corresponding selection coefficients. At random moments of time, fitness values are redrawn from the same distribution (Fig. 1a-c).

**Fig. 1.**
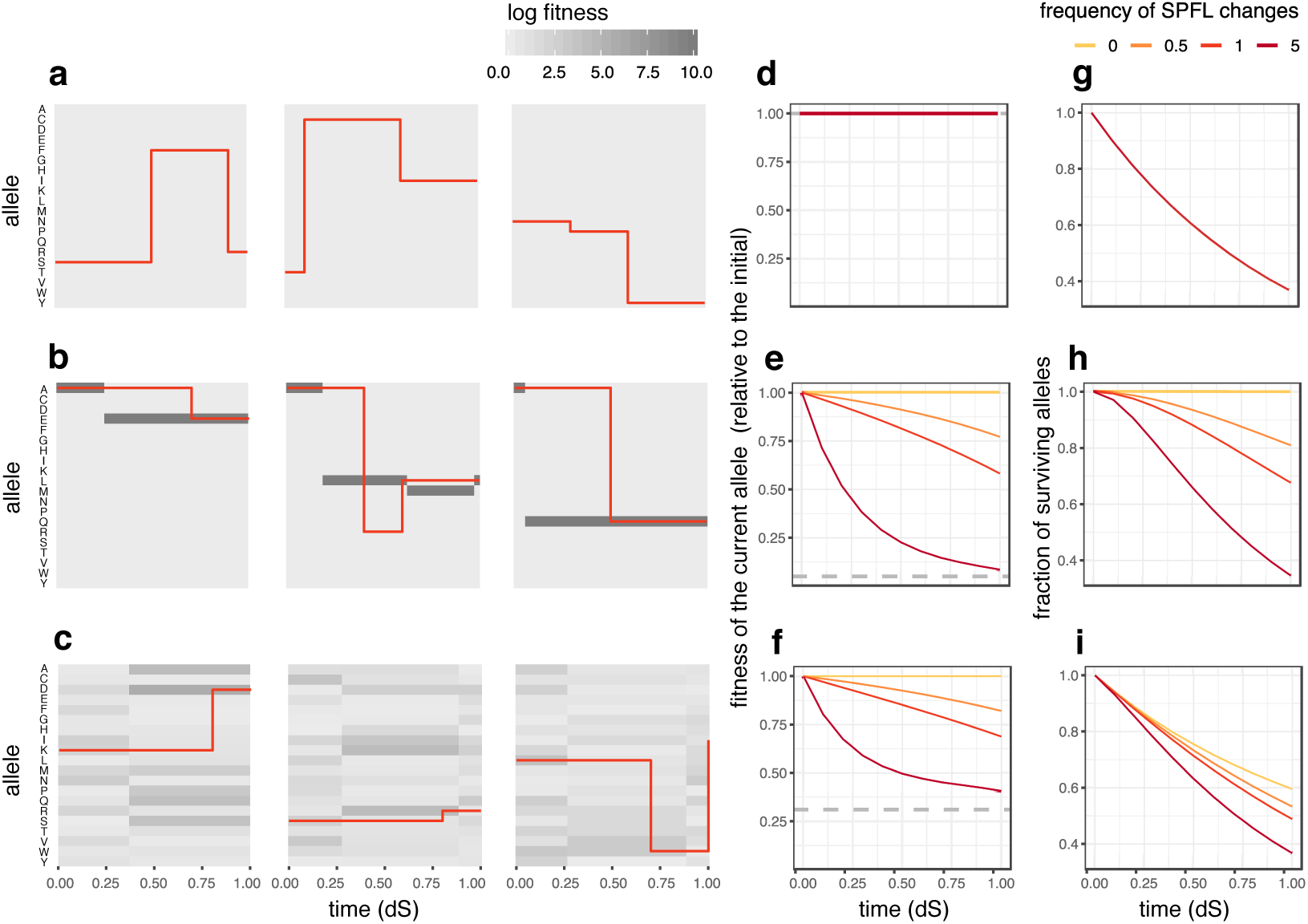
Random changes of SPFL reduce the fitness of the current allele. **a–c**, Examples of how simulated random changes in SPFLs of different shapes provoke allele substitutions. **a**, All 20 possible alleles have the same fitness (flat SPFL); **b**, One allele is substantially more beneficial than the others (rugged SPFL, log fitness vector = [10, 0, …, 0]), **c**, log fitness values are drawn from the gamma distribution with shape and rate = 1. Each panel shows the history of one simulated amino acid site; red lines represent the current allele at the site, with vertical red lines indicating substitutions. SPFL changes randomly at the average rate of one change per time required for one neutral substitution (1 dS). **d–f**, Changes in the average fitness of the current allele with evolutionary time under random SPFL changes. If the SPFL is static (frequency of SPFL changes = 0), the fitness of the current allele remains constant. The same is true for the flat landscape (**d**). Otherwise, the fitness of the current allele declines for both the rugged (**e**) and the gamma-distributed (**f**) SPFLs, i.e., the current allele undergoes senescence, and the more frequently the SPFL changes, the more rapid is this decline. The mean fitness across all possible alleles is shown with a dashed line. **g–i**, The fraction of surviving alleles as a function of time since the beginning of the simulation. For both the rugged (**h**) and the gamma-distributed (**i**) SPFLs, random changes in the landscape increase the rate at which the original allele is lost (**h–i**), unless the SPFL is flat (**g**). For **d–i**, 95% confidence bands based on 10 repeats are plotted (but too narrow to be seen).

As a result of selection, the fitness of the current allele is on average higher than that of the other ones (Fig. 1b-c); in particular, if selection is strong, the site is typically occupied by the best-possible allele (Fig. 1b). However, as the landscape changes randomly, the fitness of this original allele, on average, decreases with time, gradually approaching the mean fitness across all possible variants (Fig. 1e,f). We term this process senescence^41^. This effect is more pronounced for the rugged landscape, when one allele is highly more beneficial than the others (Fig. 1e), and less pronounced when selection is weaker (Fig. 1f).

The decline of the current allele fitness due to fluctuating selection leads to an increase in the rate at which it is lost (Fig. 1h,i), in line with the quenched theory of fluctuating selection^34,37^, Supplementary Text 2).

### Senescence and entrenchment result in opposite substitution patterns

Therefore, the two different modes of change of the SPFL are expected to produce the opposite dynamics of the fitness of the allele that currently occupies the site. If the current allele is favored by epistatic interactions with other sites, it will be entrenched, i.e. its fitness, compared to that of other alleles at this site, is expected to increase with time. By contrast, random SPFL changes that occur without regard to the identity of the allele currently occupying the site are expected to decrease the fitness of the current allele with time, leading to senescence.

This dichotomy can, in principle, be used to distinguish between these two modes of SPFL changes. To infer the changes in the relative fitness of an allele with time, we study the differences in the rate at which it is lost in the course of evolution between different groups of species. Indeed, the relative fitness of an allele specifies the probability with which it is substituted by another allele per unit time in the course of evolution^42^.

Let us assume that a substitution of an ancestral variant A for another variant B (allele gain) has occurred at some internal branch of the phylogenetic tree, and this current allele B has remained fixed in several extant species (Fig. 2a). Following the gain of B, its losses could occur, for example, as a result of a reversal to A or a substitution to some other allele C. If the SPFL for this site has remained static (the fitness of the current allele B has not changed, Δf_B_= 0), the probability of replacement of B is independent of time elapsed since its gain.

**Fig. 2.**
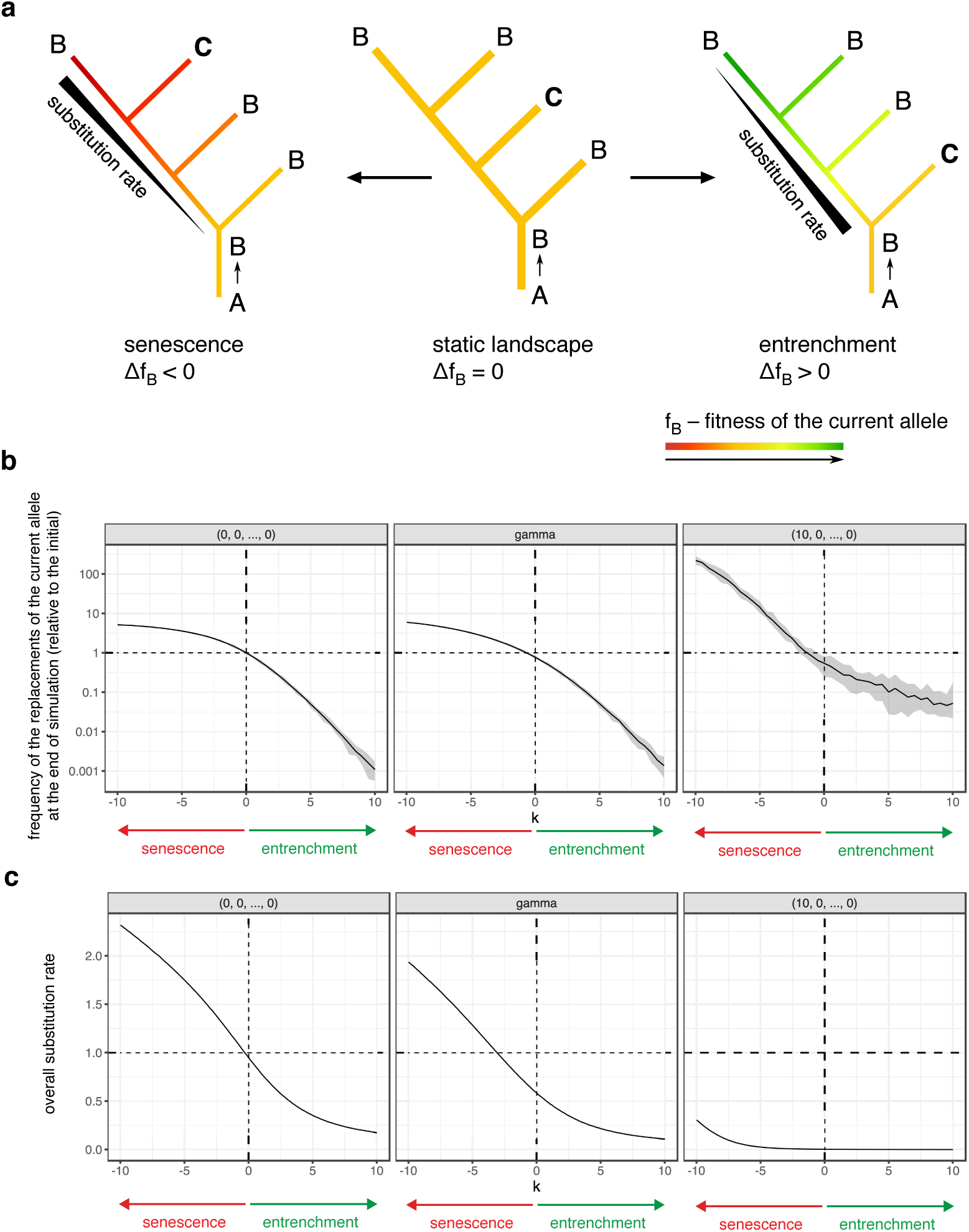
Replacement patterns of the current allele reflect changes of its fitness. **a**, On the static landscape (if the fitness of the current allele B remains constant, Δ *f*_*B*_ = 0), the probability that B is replaced per unit time does not depend on the time since its gain A *→* B. Under entrenchment (right), B becomes more favourable with time (Δ *f*_*B*_ > 0); therefore, the A *→* B substitution rate declines and there are fewer substitutions observed on “late” branches of the phylogeny. Alternatively, under senescence (left), the fitness of B decreases (Δ *f*_*B*_ < 0), leading to an increase in the rate of its loss with time. **b**, The dynamics of replacements of the current allele changes in its fitness. SPFLs of different shapes were simulated: flat SPFL (log fitness vector = [0, 0, …, 0]), rugged SPFL (log fitness vector = [10, 0, …, 0]), and gamma-distributed SPFL. The log fitness of the current allele was linearly changing with time at rate *k*. If there are no changes in the fitness of the current allele (*k* = 0), the frequency of substitutions on the “early” branches is similar to that on the “late” branches. If the current allele is senescing (*k* < 0), more substitutions are observed on the “late” branches; on the contrary, under entrenchment (*k* > 0), there are more substitutions on the “early” branches. **c**, Changes in the current allele fitness also affect the overall substitution rates: senescence leads to an increase in the rate of substitutions as compared to a static SPFL of the same shape, while entrenchment reduces the substitution rate. For **b**,**c**, 95% confidence bands based on 10 repeats are shown.

By contrast, under senescence, the fitness conferred by B decreases with its age (Δf_B_< 0), and the probability of its replacement increases with it. In this case, we will observe a higher rate of substitutions on the branches originating much later than the allele gain, compared to the branches leading to close descendants (Fig. 2a, left). Finally, under entrenchment, the fitness of the current allele increases (Δf_B_> 0), so the frequency with which B is lost declines with its age (Fig. 2a, right).

To test the validity of this approach, we used SELVa^43^ to simulate molecular evolution at individual sites assuming that the fitness of the allele currently residing at the site changes with time. Specifically, we assume that the log fitness of the current allele is initially drawn from a predefined distribution, and then changes with time linearly with rate *k*. Positive values of *k* correspond to an increase in the fitness of the current allele, i.e., entrenchment, while negative values correspond to a decrease in its fitness, i.e., senescence (see Methods). As expected, entrenchment (*k* > 0) results in a high rate of substitutions immediately after the ancestral substitution, but a reduced rate later on. By contrast, under senescence (*k* < 0), the rate of substitutions increases with time since the allele gain (Fig. 2b). An alternative mode of simulation of senescence, whereby random changes in SPFL and the molecular evolution caused by them are modeled explicitly, gives the same results (Fig. S1).

Besides the phylogenetic distribution of substitutions, the mode and rate of SPFL change also affect the overall rate of molecular evolution (Fig. 2c). Compared to a static landscape of the same shape, entrenchment reduces the substitution rate, as the time-averaged fitness of the current allele is higher, and therefore its substitutions rarer. Conversely, under senescence, many of the substitutions of the current allele are advantageous, increasing the overall rate of evolution. If selection is strong, the overall substitution rate is low, as the site is usually occupied by the optimal allele; nevertheless, this rate is increased by senescence, whereby the identity of the optimal allele changes occasionally.

### Senescence and entrenchment at single-allele resolution

Large phylogenies allow detecting changes in substitution frequencies for specific alleles. Each originating allele, e.g. an amino acid arising at an individual site from an ancestral amino acid substitution, can be inherited by multiple descendant lineages leading to different extant species. Ancestral state reconstruction can then be used to infer the lineages at which this allele has been lost due to a reversion or substitution to a different amino acid. If enough such lineages are available, this allows to trace the decline or increase in the rate of allele substitution since its origin, i.e., entrenchment or senescence.

We applied binomial logistic regression to detect changes in substitution frequencies with the age of the current allele along the phylogeny for five mitochondrial genes of Metazoa^14^. The regression was performed separately for each allele B with a known time of origin (corresponding to allele gain A → B) at each site. Among the 42,637 such alleles, we identified 28 alleles for which the frequency of replacement significantly increased with time since their origin (i.e. senescing alleles), and 21 alleles where it decreased (i. e. entrenched alleles) at 5% false discovery rate (Fig. 3a, Table S1). The examples of phylogenies indicating allele replacements at senescing and entrenched alleles are shown in Fig. 3bc.

**Fig. 3.**
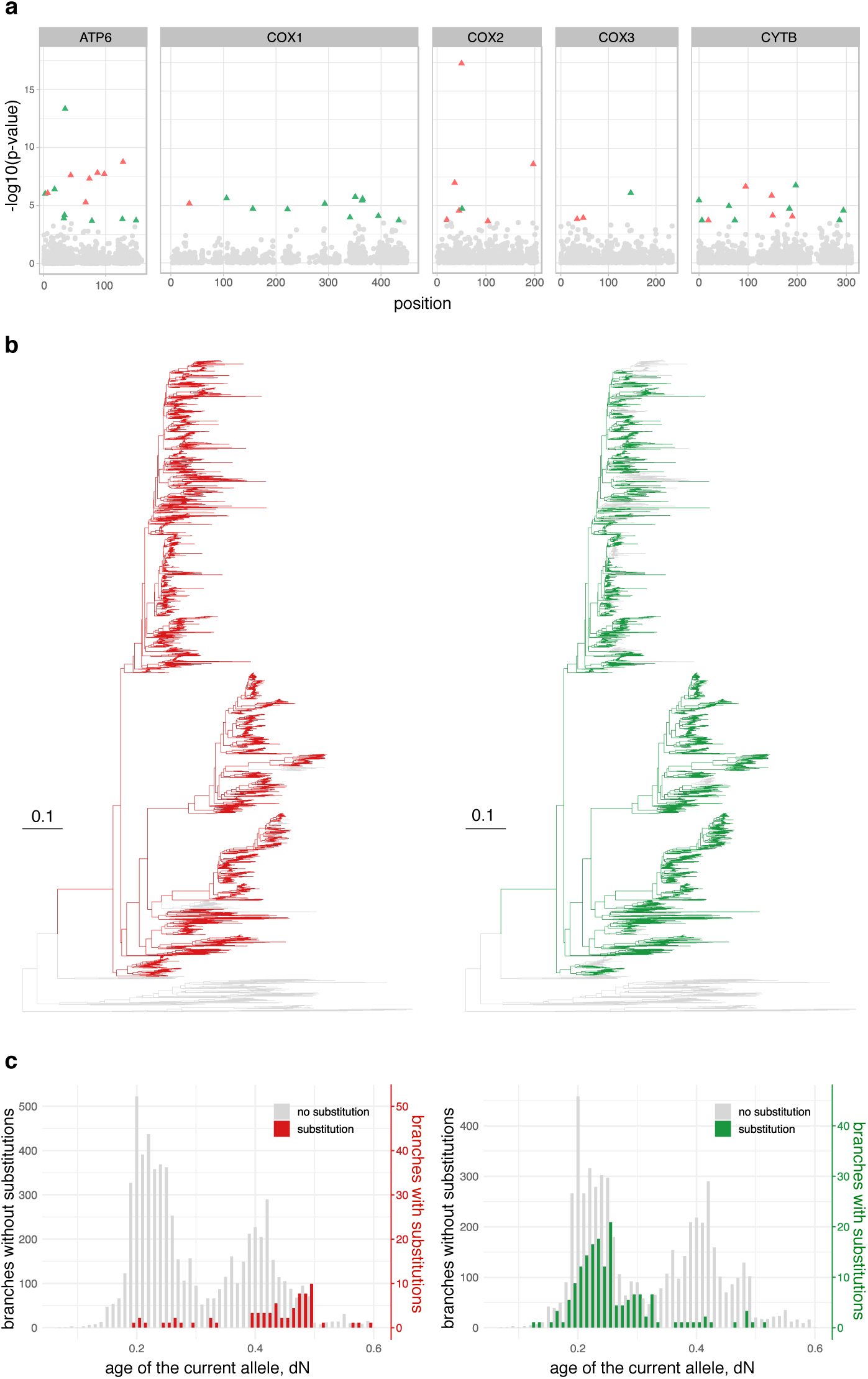
Senescence and entrenchment of individual alleles in the mitochondrial genes of Metazoa. **a**, Manhattan plot of senescing and entrenched alleles. Only the alleles with a known phylogenetic position of origin, i.e., those that were not yet present in the tree root, were analyzed; a single genomic site can contain zero, one or several alleles. P-values are calculated using binomial logistic regression. The alleles demonstrating significant senescence under 5% FDR are shown in red; the alleles demonstrating entrenchment are shown in green. No amino acid sites contained more than one significantly senescing or entrenched alleles. The list of significantly senescing or entrenched alleles is shown in Table S1. **b**, Examples of senescing (COX2 position 56, red) and entrenched (ATP6 position 71, green) alleles. The contiguous segment of the phylogeny carrying the derived allele is shown in color. **c**, Distribution of substitutions along the lifetime of alleles shown in (**b**). For the senescing allele, the phylogenetic branches corresponding to allele replacements (red) originate later than the branches without replacements (gray). Conversely, the entrenched allele is more frequently replaced right after its origin (green).

### Heterogeneity of alleles leads to an artifactual signal of entrenchment

While phylogenies spanning hundreds and thousands of species, like those available for mitochondrial proteins, allow to measure the changes in the substitution rate for individual alleles, in smaller phylogenies, the number of substitutions experienced by an allele can be insufficient for such an analysis. Still, it may be possible to identify the prevailing patterns of substitutions by pooling alleles together. However, such pooling can be problematic: even in the absence of SPFL changes, the rate of substitution can appear to change with time since allele origin if alleles with different time-invariant substitution rates are pooled together, confounding inference of SPFL changes.

Indeed, consider a set of alleles, each characterized by its own substitution rate that is stationary (constant in time) but differs between alleles. While the replacement rate may be constant for each allele, so that the time to replacement is characterized by an exponential distribution, it will not, in general, be exponentially distributed in the resulting heterogeneous dataset. Instead, the frequency of substitution will appear to decline with time (Fig. 4a), making it non-stationary and mimicking entrenchment of the current allele. The problem of data heterogeneity leading to decreasing hazard function is well known in demographic inference^44,45^, and has been previously appreciated in the inference of substitution rates dynamics in molecular evolution^28,46^. Notably, no mixture of stationary processes can give rise to an increase in the substitution rate, i.e., senescence^44^.

**Fig. 4.**
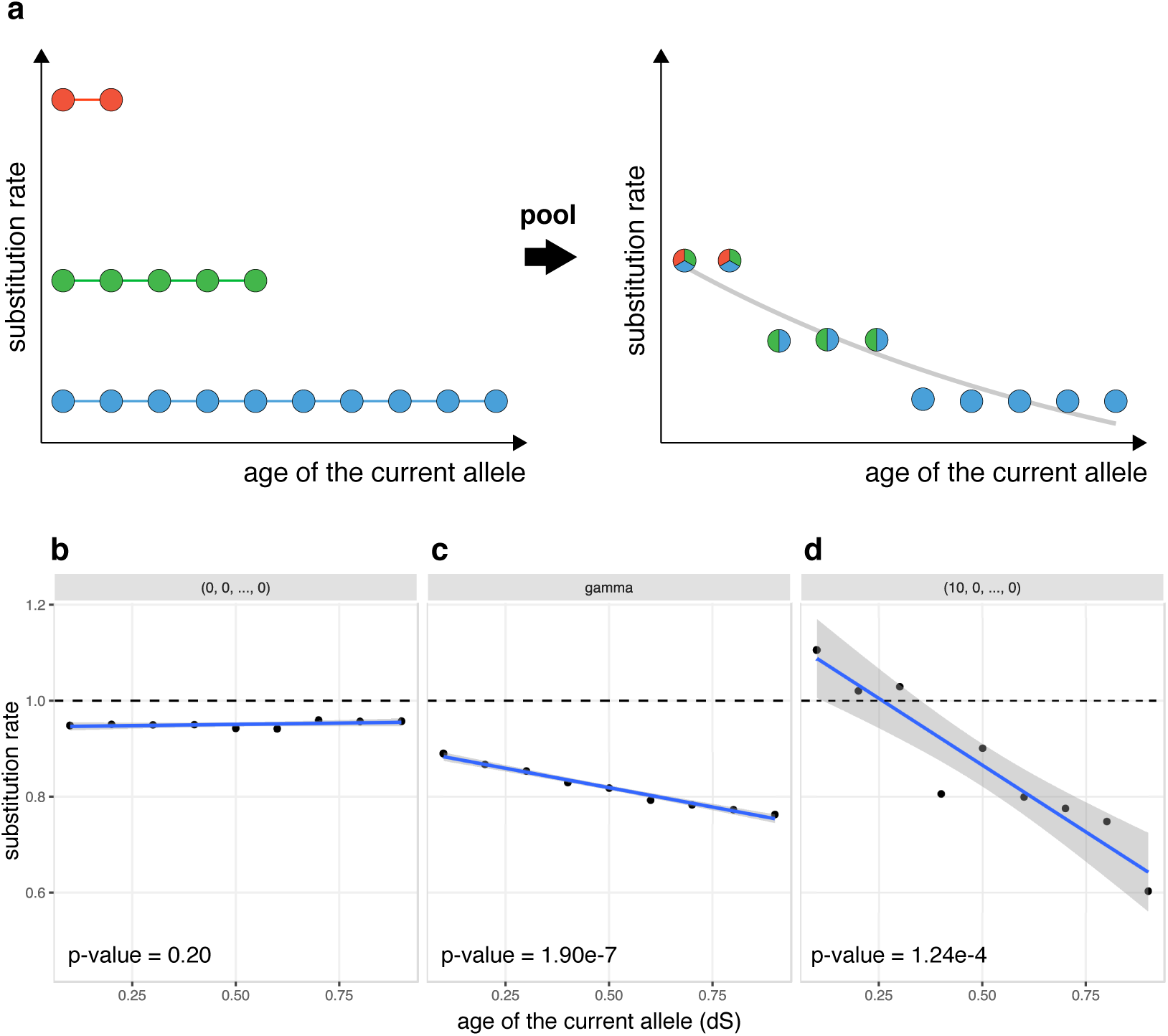
Heterogeneity of substitution rates among alleles on static SPFLs imitates entrenchment. **a**, Consider three classes of alleles, characterized by different constant substitution rates: fast (red), moderate (green), and slow (blue) alleles. For each allele, we can calculate the substitution frequency on the phylogenetic branches located at different evolutionary distances from the gain of that allele (each dot corresponds to a single branch). If the dynamics of replacement of these alleles are analyzed separately for each substitution-rate class, no spurious signal of entrenchment or senescence is observed (left). However, if alleles from different classes are pooled together, the substitution frequency appears to decrease with time, mimicking the signal of entrenchment (right). **b**, On a static flat SPFL (log fitness vector = [0, 0, …, 0]), i.e., when the substitution rates for all 20 alleles are the same, the frequency of replacement of the current allele on some branch of the phylogenetic tree does not depend on the time since the allele gain (each dot corresponds to a single branch). However, on a gamma-distributed SPFL, the heterogeneity of fitness of the alleles produces entrenchment-like decline of replacement rate of the current allele with time, although the SPFL remains static. The effect is even more pronounced on a more rugged SPFL (log fitness vector = [10, 0, …, 0]).

It is obvious that heterogeneity of substitution rates arises from pooling of different amino acid sites. More subtly, it also arises within individual sites as a result of differences between rates of substitution of different alleles. Substitution rate is the product of mutation and fixation rates, and this heterogeneity will arise due to any differences in either of these factors between alleles. For example, consider a single site which is non-neutral, i.e., such that different alleles confer different fitness. Such alleles will be characterized by different replacement rates (lower for high-fitness alleles, and higher for low-fitness alleles), and pooling over different alleles over the course of evolution of this site (or, identically, over different independent and identically distributed sites) would lead to heterogeneity of substitution rates and to an apparent decline in substitution rates with time since allele origin.

To show this, we simulate molecular evolution on static SPFLs of different shapes. If all alleles have the same fitness, i.e., if all substitutions are neutral (“flat” landscape), the substitution rate is independent of time since allele origin (Fig. 4b). By contrast, if the fitness values of alleles are drawn from a gamma distribution, so that different alleles have different fitness, the substitution frequency decreases with the age of the current allele, even though the SPFL doesn’t change (Fig. 4c). On a more rugged SPFL, when one allele is much more fit than all others, this decline is even sharper (Fig. 4d).

### Inferring senescence and entrenchment from phylogenetic distribution of substitutions

To address the problem of heterogeneity of alleles, we reconstructed the dynamics of the fitness landscape while accounting for differences between alleles in mutation rates and “baseline” selection. In the absence of prior information about the distribution of these characteristics, it was impossible to reconstruct the explicit likelihood function for the substitution rates. Instead, we used the approximate Bayesian computation (ABC)^47^ approach to obtain the posterior distribution of the rate of current allele fitness change per unit time (*k*). ABC depends on a set of summary statistics to evaluate the difference between the simulation results and the data. We used two summary statistics, each aggregating over all individual alleles, which reflect the age-dependent dynamics of substitution rates (see Methods, Fig. S2).

We used two models for parameter inference. Under the two-parameter model, we assumed that log fitness values for individual alleles were drawn from a gamma distribution with rate and shape parameters denoted by *alpha*, and the log fitness of the current allele at all sites changed linearly with rate *k*. Under the three-parameter model, the fitness changed linearly only for a fraction of alleles, while the fitness of the remaining alleles was invariant (see Methods for details).

Simulations show that both models perform well in identifying senescence and entrenchment under a broad range of parameters, and are robust to overall substitution rate, phylogeny shape, pooling of sites with diverse characteristics and errors in ancestral state reconstruction (see Methods, Table S2, Fig. S3-7).

### Positively selected sites show strong senescence

We applied the developed ABC approach to protein sequences of vertebrates and insects (Fig. S8). To understand how the direction of fitness change depends on the overall conservation of an amino acid site, in both datasets, we classified all codon sites by the type of selection acting at the site, on the basis of the ratio of nonsynonymous and synonymous substitutions per site (ω): negatively selected sites (ω < 1), neutral (ω = 1) or positively selected (ω > 1), and analyzed them independently (Table 1). The three-parameter model provided a better fit to the data than the two-parameter model (Fig. S9-10), so we used the former model for the analysis.

**Table 1.** The analyzed datasets, the corresponding estimates of the rate of entrenchment (*k* > 0) or senescence (*k* < 0) and the fraction of alleles that experience these processes. The values show the median of the ABC posterior distribution of parameter values; numbers in parentheses represent the 95% posterior probability intervals.

|  | # of sites |  | # of alleles |  | k | fraction of senescing<br>or entrenched alleles |
| --- | --- | --- | --- | --- | --- | --- |
| Vertebrates |  |  |  |  |  |  |
| $\omega < 1$ | 2 598 701 | (87.8%) | 574 137 | (36.4%) | 62.3 (25.1, 100.0) | 0.99 (0.95, 1.00) |
| $\omega = 1$ | 348 047 | (11.8%) | 917 340 | (58.1%) | 45.3 (-36.8, 98.9) | 0.42 (0.15, 0.90) |
| $\omega > 1$ | 13 189 | (0.4%) | 86 852 | (5.5%) | -23.3 (-94.7, -3.5) | 0.18 (0.0, 0.86) |
| **Total** | **2 959 937** | **(100%)** | **1 578 329** | **(100%)** |  |  |
| Insects |  |  |  |  |  |  |
| $\omega < 1$ | 2 699 432 | (93.2%) | 429 800 | (48.8%) | 47.8 (-1.1, 87.9) | 0.81 (0.47, 1.00) |
| $\omega = 1$ | 185 829 | (6.4%) | 413 101 | (46.8%) | 9.4 (-80.2, 91.4) | 0.16 (0.01, 0.92) |
| $\omega > 1$ | 9 698 | (4.4%) | 38 577 | (4.4%) | -22.6 (-88.7, 52.0) | 0.25 (0.03, 0.92) |
| **Total** | **2 894 959** | **(100%)** | **881 478** | **(100%)** |  |  |

In vertebrates, both the fraction of senescing or entrenched alleles and the value of *k* for them depended on the mode of selection acting at the site. The 36% of alleles originating at negatively selected (ω < 1) sites demonstrated strong evidence for entrenchment: we estimate that all of them are entrenched, indicating that the fitness of the current variant increases with time since its origin (Fig. 5a; Table 1). By contrast, of the 6% of alleles arising at positively selected sites (ω > 1), 18% experience senescence (Fig. 5c), indicating a decrease in the fitness of the current allele. While we are unable to distinguish robustly between a low fraction of alleles undergoing strong senescence and a high fraction of alleles undergoing weak senescence (Fig. 5c), the 95% posterior probability interval does not include *k=*0, rejecting stationarity. The neutral sites demonstrate an intermediate signal with little evidence for entrenchment or senescence (Fig. 5b). A similar pattern is observed in phylogenies of insects (Fig. 5d-f).

**Fig. 5.**
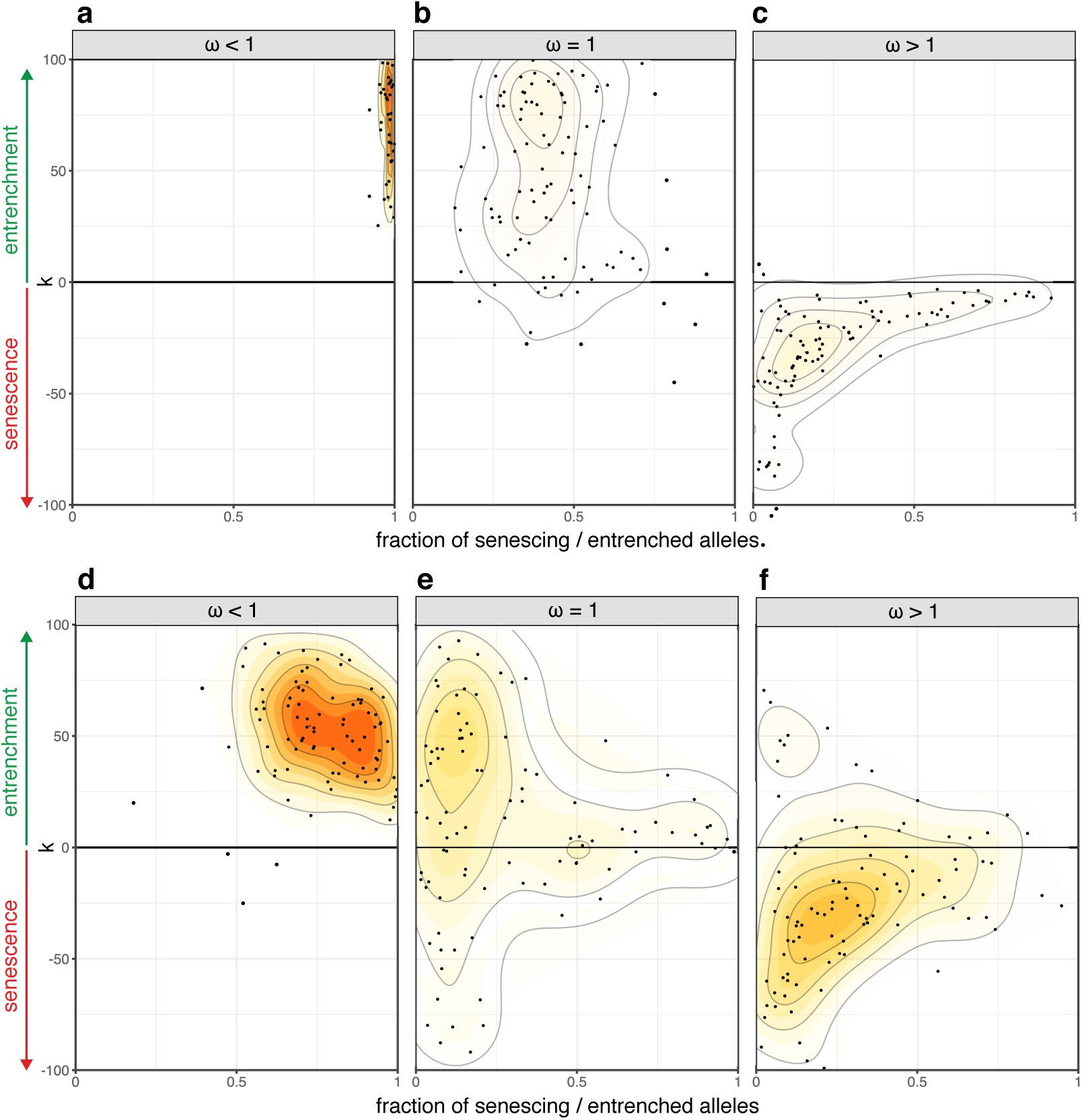
ABC estimates of the rate of senescence or entrenchment *k* and the fraction of alleles with changing fitness for protein sequences of vertebrates and insects. For each dataset, the posterior distribution of parameters under 1% acceptance threshold after local Ridge regression adjustment is shown. Positive values of *k* correspond to entrenchment, while negative values of *k* correspond to senescence of the current allele. In vertebrates, sites under negative selection show strong entrenchment (**a**), neutral sites demonstrate the intermediate signal (**b**), and positively selected sites are senescing (**c**). A similar pattern in observed in insects (**d-f**).

## Discussion

While the direction of changes in fitness in the course of evolution is unpredictable for an individual allele, there are certain statistical regularities. Previous work has shown that, at a site involved in epistatic interactions with other sites, the relative fitness conferred by the incumbent allele is expected to increase with time since its origin^9,18,22,23^. Acting alone, i.e., if the overall fitness landscape is static, this process of entrenchment should make the propensities existing at individual sites more pronounced. This, in turn, should limit the level of divergence between highly divergent sequences, although reaching this level may take a very long time^9,10,48–51^.

Here, we consider the dynamics of allele fitness due to changes in the overall fitness landscape itself. We show that, if the direction of these changes is independent of the current position of the population in the genotype space, the expected mean dynamics — senescence — is opposite to that of entrenchment. We design a method to distinguish the two patterns from the phylogenetic distribution of substitutions and find that entrenchment is prevalent at negatively selected sites, while senescence is ubiquitous at sites undergoing adaptive evolution under positive selection. This is consistent with the status of senescence as another facet of positive selection and hints at potential approaches for its detection.

We have previously revealed the acceleration of substitutions over the course of allele lifetime in the evolution of influenza A virus^41^. This pattern has been mainly observed at sites associated with avoidance of the host immune system pressure, and we interpreted it as evidence for negative frequency-dependent selection actively disfavoring the current allele^41^. Here, we show that this type of selection is not a prerequisite for senescence. Instead, senescence is expected whenever selection changes without regard to the identity of the current allele. The genome-wide signal of senescence observed in positively selected sites of vertebrate and insect genomes is perhaps more consistent with the latter scenario.

## Methods

### Data

We used multiple alignments of exons of vertebrates and insects from the UCSC Genome Browser database together with the corresponding phylogenies (Fig. S8)^53^. Columns with gaps were excluded. From these alignments, we reconstructed the alleles in the internal nodes of phylogenetic trees with *codeml*^54^. We re-estimated the lengths of individual branches as the average frequency of amino acid substitutions per site on this branch. We classified codon sites as negatively selected (ω < 1), neutral (ω = 1) or positively selected (ω > 1) using Bayes empirical Bayes (BEB) method as implemented in the *PAML* package^55^. The size of the datasets and the number of sites and substitution subtrees in each bin are shown in Table 1. The mitochondrial dataset consists of the amino acid alignment of five proteins for several thousand metazoan species^14^.

### Simulations

Simulations of amino acid sequence evolution were performed with the SELVa simulator^43^. SELVa is a forward-time Markov chain simulator that allows the user to model sequence evolution along a predefined phylogenetic tree on static or dynamic SPFLs. The user can specify both the shape of SPFL (i.e., the vector of allele fitnesses for a single position in the genome) and the rule for its change. In this work, we used three types of SPFLs for amino acid sites with 20 possible alleles: flat SPFL corresponding to neutral sites (no substitution leads to change of fitness, log fitness vector is (0, 0, …, 0)); rugged SPFL (one allele is highly preferable over the other ones, log fitness vector is (10, 0, …, 0)) and gamma-distributed SPFL, where the log fitness values for alleles are randomly chosen from the gamma distribution with user-defined parameters.

We used two modes of SPFL change. In the random change mode, the SPFL changes are a Poisson process with a user-defined rate. In this case, fitness values are either reshuffled between alleles (for the flat or rugged SPFL) or redrawn from the same distribution (for gamma-distributed fitnesses). In the current allele-dependent mode, the log fitness of the current allele increases or decreases linearly with time. The user can define the rate at which the fitness of the current allele changes per unit time (*k*) and the length of the fixed time interval between changes (*Δt*). The new log fitness of the current allele *B* is defined as

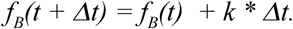

The fitnesses of other alleles remain unchanged. Positive values of *k* correspond to entrenchment of the current allele, which means its fitness increases with time, and negative ones, to senescence, so that its fitness decreases. If a substitution occurs, i.e. the current allele is replaced with another one, the fitness of the new allele changes with the rate *k*. In this work, we used *Δt* = 0.01 dS, which is small enough to simulate the gradual change of current allele fitness.

The substitution rates are scaled to the neutral substitution rate. Individual sites are simulated independently, and interactions between them are not modeled directly. We also do not model the overall substitution rate per site explicitly in the simulations; the heterogeneity of substitution rates among sites arises from the randomly chosen gamma-distributed alleles fitness values.

To demonstrate the effects of randomly changing SPFLs, patterns of substitutions emerging due to senescence and entrenchment, and the influence of allele heterogeneity, we used the simple model phylogenetic tree (Fig. S12). We used phylogenies of vertebrates and insects for ABC-based inference of the dynamics of the current allele fitness in the evolution of their protein sequences (Fig. S8).

### Substitution subtrees

For every replacement A → B on any internal branch of the phylogeny, we can define the corresponding substitution subtree, namely, the contiguous segment of the phylogeny where every internal node carries the derived variant B. For this subtree, B is the current allele, and the initial substitution A → B is the allele gain (Fig. S2). Within a substitution subtree, the current allele B can be replaced by the ancestral variant A or some other variant C in the course of allele loss(es).

A genomic position can carry no substitution subtrees if it is fully conservative, or carry one or more substitution subtrees in a number equal to the number of substitutions on the internal phylogenetic branches in this position. This means that rapidly evolving sites possess more substitution subtrees than conservative ones.

We intend to define statistics that represent the conditional probability of the allele B to be replaced at a single branch, given that B was occupying the genomic position at the moment. This probability is completely defined by the alleles fitness vector at this point of time (that is, the SPFL) and the mutation rate, so it can be used to detect changes of SPFL within the substitution subtree. The substitution subtrees perspective allows us to estimate this conditional probability as a function of the age of the current allele B, i.e. the evolutionary time since it was gained (with its gain defining the root of the subtree).

### Extraction of the signal of senescence or entrenchment for groups of alleles

An individual substitution subtree usually does not provide enough data to make conclusions about SPFL changes. Therefore, we have to pool together replacement data from different subtrees and sites to identify SPFL changes with confidence. However, pooling data from different subtrees creates a signal of pseudo-entrenchment due to heterogeneity of evolution rate and unevenness of SPFLs^28,46^. One approach to adjust for these confounding factors is to estimate the mean evolution rate of a subtree, which combines both the mutation rate of the site and the effect of the fitness variance of the current allele, and to use this value in the model fitting, e.g. in the maximum likelihood (ML) framework. However, our phylogenies are not deep enough to perform ML estimates: the number of substitutions per subtree is too low, while the variance of branch lengths is too large. Instead, to account for the confounding effects, we used the approximate Bayesian computation approach (ABC).

Since the age-dependent patterns of substitutions are sensitive to data heterogeneity and the shape of SPFLs, we can’t directly measure the rates of current allele fitness change. To estimate the strength and abundance of senescence and entrenchment in the evolution of protein sequences we used rejection ABC (approximate bayesian computation) with Ridge regression adjustment as implemented in the *abc* package for R^56^. ABC is a popular method for parameter inference if the likelihood function is not known.

Simulations for ABC were also performed by the SELVa simulator. Importantly, simulations were performed not for the full phylogenetic trees, but for individual substitution subtrees. For each dataset of interest, we extracted the list of subtrees generated by substitutions in this dataset. Then SELVa was launched with given parameters based on each subtree, and the number of sites in the simulation was equal to the number of appearances of this subtree. The results were merged across subtrees, and summary statistics were calculated. This approach has advantages in comparison to simulations based on full phylogenetic trees. First of all, our summary statistics are based on subtrees only, and the list of substitution subtrees is more indicative of the dataset than simply the number of sites: we don’t have to wait until the ancestral substitution occurs, but begin the simulation at the same moment when this substitution happens. Secondly, since the subtrees are smaller, the simulation runs faster.

We used two model functions for ABC. The first one is based on the assumption that all sites in the dataset are susceptible to senescence or entrenchment of the same strength (two-parameter model). It requires two parameters: *alpha* rate parameter of gamma distribution of fitness values of alleles (see Simulations) and the rate of the current allele’s fitness change *k*.

The second model represents a mixture of sites with static SPFL (*k* = 0) and sites under senescence (*k* < 0) or entrenchment (*k* > 0). It takes three parameters as input: the parameter of gamma fitness distribution *alpha*, the rate of the SPFL change *k* and the fraction of substitution subtrees (which corresponds to the fraction of alleles) under senescence or entrenchment with the rate *k*. The simulated values for *k* were distributed uniformly from −100 to 100, the fraction of alleles under senescence or entrenchment was also distributed uniformly from 0% to 100%, and *alpha* was distributed log-uniformly from −1.5 to 1.

The number of amino acid sites in the datasets with different site-specific ω varies from 10^3^ to 10^6^, and so varies the number of substitution subtrees in these datasets (Table 1). To take into account the variance of summary statistics for smaller datasets, the ABC model function takes a list of subtrees and their count in the given dataset as input and generates simulations of the same size (but not larger than 100 000 substitution subtrees due to runtime restrictions). For all datasets, we used Ridge regression algorithm for parameter estimation as implemented in the *abc* package.

### Summary statistics

After evaluating a range of possible summary statistics for ABC, we ended up using two statistics based on the dynamics of allele replacement. All branches across all subtrees in the simulation are pooled together and used to calculate the following linear regression:

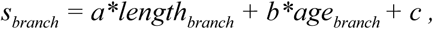

where *s*_*branch*_ is the frequency at which the current allele was lost lost on the given branch, *length*_*branch*_ is the average length of this branch across all substitution subtrees, and *age*_*branch*_ is the age of the current allele, i.e. the distance from the root of the substitution subtree to the branch. As summary statistics, we used the values of *a* and *b*. Fig. S11 shows how *a* and *b* depend on the simulation parameters: they strongly depend on the direction and the rate of current allele fitness change *k* and the fraction of alleles susceptible to these changes.

### ABC validation

Since our method is based on time-dependent dynamics of substitutions, its efficiency is expected to depend on the depth of the phylogenetic tree: short trees won’t provide the timescale sufficient to detect changes of the current allele substitution rate. We examined the accuracy of the two-parameter model using test phylogenies with simple topology and internal edges of varying lengths (Fig. S3a). Indeed, both the prediction error and the width of the 95% posterior probability interval are the smallest for the phylogenies with internal branches of intermediate lengths (Fig. S3cd). While the shortest trees don’t provide the time range sufficient to detect SPFL changes, the largest ones lack resolution and also result in bigger prediction error.

We validated the ABC pipeline for parameter inference using SELVa simulations based on the reconstructed phylogeny of 53 vertebrates and the *abc* package for R^56^. To cross-validate ABC performance under different tolerance rates and to evaluate the accuracy of parameter estimation for both two-parameter and three-parameter models, we calculated prediction error for parameters based on 100 randomly chosen simulations with the cross-validation function of the *abc* package. The prior size was 10^4^ simulations for both models. The cross-validation results are shown in Table S3 and Fig. S4-5. The tolerance level 0.01 was chosen for both models. Cross-validation tests for parameter inference with our ABC pipeline show that we can accurately estimate the parameters of both models using the selected set of summary statistics.

Next, we asked whether our method is sensitive to changes in the overall evolution rate. For each model, we generated the testing set of 100 simulations with randomly chosen parameters with normal and twofold increased substitution rate and then used ABC to infer the parameters. We demonstrate that, although the magnitude of *k* was overestimated for simulations with accelerated evolution rate, the estimates were not biased in any direction (p-value=0.53, Fig. S6-7, top panel; Table S2). However, the fraction of senescing or entrenched alleles in the three-parameter model was overestimated for simulations with accelerated evolution rate, but the magnitude of the bias was not large (on average 0.10, p-value=3e-10).

We also checked whether out method allows to confidently distinguish between senescence and entrenchment. The confusion matrices based on the same testing set of simulations show that the frequency of errors is 0% for the two-parameter model and 1% for the three-parameter model, and concern only simulations with the small magnitude of *k* (< 1) (Table S2). In the few erroneously classified cases, the 95% probability interval for *k* also overlapped with zero.

SELVa simulator stores the sequences of internal nodes of the phylogenetic tree (the ancestral sequences) so that the history of simulated amino acid replacements is known exactly. However, for real data, we use *codeml* to reconstruct the ancestral sequences, and this reconstruction can be erroneous. To make sure that the ancestral state reconstruction does not affect the accuracy of parameter estimation, we reconstructed the ancestral sequences generated by SELVa on the basis of the sequences of terminal nodes in the same way as it was done for the actual data, and used ABC to estimate the parameters using the same procedure as above (Fig. S6-7, bottom panel). We found that ancestral states reconstruction slightly biased both *k* (by ∼3.8, p-value = 6e-4) and the fraction of alleles with changing fitness (by ∼0.08, p-value < 2e-16) upwards, but the confusion frequency remained low (0% for the two-parameter model, 0.5% for the three-parameter model), and the only erroneously classified simulation had a small fraction of entrenched alleles (0.09).

## Acknowledgments

We thank Dmitrii Ivankov and Alexey Kondrashov for useful comments on drafts of this article.

## Author information

### Contributions

A.V.S., E.N., V.V.P., A.V.F., A.D.N., and G.A.B. designed research; A.V.S., E.N. and V.V.P. performed research; A.V.S., and A.V.P. analyzed data; and A.V.S., A.V.F.,and G.A.B. wrote the paper.

