## Supplementary Text for "Senescence and entrenchment in evolution of amino acid sites"

### Supplementary Text 1

### Formal proof that random changes in SPFL decrease the fitness of the current allele

Consider a locus under selection with $N$ different alleles. These alleles differ in fitness that they confer, so that allele $i$ confers fitness $w_i$. The vector of these fitness values, SPFL, can change instantaneously. At the time of such a change, each allele is, in general, ascribed a new fitness value $w_i'$, giving rise to a new SPFL. We assume that the effective population size is constant, so that the substitution rate between alleles $i$ and $j$ depends on the difference in their fitness $s_{ij} = w_i - w_j$.

### Notation

| | | |
|---|---|---|
| $\tau_M$ | = | average time between substitutions |
| $\tau_L$ | = | average time between SPFL changes |
| $N$ | = | number of possible alleles |
| $a_i$ | = | probability that the considered position is occupied by the allele $i$ |
| $w_i$ | = | fitness of allele $i$ at the considered position before the SPFL change |
| $w_i'$ | = | fitness of allele $i$ at the considered position after the SPFL change |
| $s_{ij} = w_i - w_j$ | = | selection coefficient in favor of the allele $i$ compared to the allele $j$ at the considered position |
| $f_{ij} = f(s_{ij})$ | = | probability of $i$ to $j$ substitution in a given position as a function of the selection coefficient. |

Here we consider allele substitutions as a Markov process, and don't account for changes in allele frequencies within the population. The probability of an allele to be fixed monotonically increases with its fitness [1], so we assume that $f(s_{ij})$ is (strictly) monotonically increasing.

The sum of allele frequencies equals 1:

$$\sum_i^N a_i = 1\,. \tag{1}$$

The fitness vector is normalized so that the sum of fitness values for all alleles also equals 1:

$$\sum_i^N w_i = 1\,. \tag{2}$$

A change in SPFL may increase the fitness of the allele currently occupying an amino acid position, decrease it, or leave it unchanged. We shall prove that a random SPFL change will **on average** decrease the fitness of the current allele:

$$\beta = \langle \Delta w \rangle_{i,w'} = \langle w_i' - w_i \rangle_{i,w'} = \langle \sum_i (w_i' - w_i) a_i \rangle_{w'} < 0\,, \tag{3}$$

where the averaging is done over all alleles $i$ and over all possible new SPFLs (or, analogously, over all new allele fitness values $w'$).

### 1. Rare SPFL changes ($\tau_M \ll \tau_L$)

If the average time between SPFL changes is significantly larger that the average time between substitutions (i. e. $\tau_M \ll \tau_L$), the distribution of allele frequencies after a substitution reaches the equilibrium state, i.e. the allele frequencies vector is the eigenvector for the substitution probability matrix:

$$a_i = \sum_k f_{ik} a_k \,. \tag{4}$$

Since the allele distribution reaches equilibrium after each SPFL change, the difference between frequencies of any two alleles depends only on the fitness vector:

$$a_i - a_j = \sum_k f_{ik} a_k - \sum_k f_{jk} a_k = \sum_k (f_{ik} - f_{jk}) a_k = \sum_k (f(w_i - w_k) - f(w_j - w_k)) a_, . \tag{5}$$

If $w_i$ is larger than $w_j$, then $\forall k : w_i - w_k > w_j - w_k$. Since $f$ is monotonically increasing, this implies that $f(w_i - w_k) > f(w_j - w_k)$. Therefore, since $a_k$ can't be negative, the difference between $a_i$ and $a_j$ is also non-negative:

$$w_i > w_j \Rightarrow a_i > a_j \,. \tag{6}$$

Let's transform the equation 3 :

$$\beta = \langle \Delta w \rangle_{i,w'} = \langle \sum_i (w_i' - w_i) a_i \rangle_{w'} = \sum_i (\langle w_i' \rangle_{w'} - w_i) a_i \,. \tag{7}$$

#### 1.1. The new SPFL is independent of the previous SPFL

First assume that the SPFL changes are random and not "biased" towards any alleles, meaning that all alleles have the same average fitness $\alpha$ across all the possible SPFLs. Then, based on the alleles fitness values normalization (equation 2),

$$\alpha = 1/N \,. \tag{8}$$

Replacing the average new fitness $\langle w_i' \rangle_{w'}$ in eqn. 7 with the value from eqn. 8,

$$\beta = \sum_i (\langle w_i' \rangle_{w'} - w_i) a_i = \sum_i (1/N - w_i) a_i \,. \tag{9}$$

The fitness values of the alleles should remain normalized after the SPFL change:

$$\sum_i^N w_i' = 1 \,. \tag{10}$$

The fitness of the allele $i$ can be either less than $1/N$ (let's denote this set of alleles as $N^-$), greater than $1/N$ ($N^+$), or equal to $1/N$ ($N^0$). Then

$$\beta = \sum_{i \in N^+} (1/N - w_i) a_i + \sum_{i \in N^-} (1/N - w_i) a_i + \sum_{i \in N^0} (1/N - w_i) a_i = \sum_{i \in N^+} (1/N - w_i) a_i + \sum_{i \in N^-} (1/N - w_i) a_i \,. \tag{11}$$

Let's denote the allele with the smallest fitness greater than $1/N$ as $i_1$ and the allele with the largest fitness less than $1/N$ as $i_2$:

$$\begin{aligned} w_{i_1} &> 1/N, \\ \forall i \in N^+ : w_i &\geq w_{i_1} \,; \\ w_{i_2} &< 1/N, \\ \forall i \in N^- : w_i &\leq w_{i_2} \,. \end{aligned} \tag{12}$$

Then, from equation 6,

$$\begin{aligned} \forall i \in N^+ : a_i &\geq a_{i_1} \,; \\ \forall i \in N^- : a_i &\leq a_{i_2} \,; \\ a_{i_1} &> a_{i_2} \end{aligned} \tag{13}$$

$$\sum_{i\in N^+}(1/N - w_i)a_i < \sum_{i\in N^+}(1/N - w_i)a_{i_1}$$
$$\sum_{i\in N^-}(1/N - w_i)a_i < \sum_{i\in N^-}(1/N - w_i)a_{i_2}\,. \tag{14}$$

Then we can estimate $\beta$ from equation 11:

$$\beta = \sum_{i\in N^+}(1/N - w_i)a_i + \sum_{i\in N^-}(1/N - w_i)a_i < \sum_{i\in N^+}(1/N - w_i)a_{i_1} + \sum_{i\in N^-}(1/N - w_i)a_{i_2} =$$
$$a_{i_1}\sum_{i\in N^+}(1/N - w_i) + a_{i_2}\sum_{i\in N^-}(1/N - w_i)\,. \tag{15}$$

Since the fitness values of individual alleles are normalized,

$$\sum_{i\in N^+}(1/N - w_i) + \sum_{i\in N^-}(1/N - w_i) = \sum_i^N(1/N - w_i) = \sum_i^N w_i - N(1/N) = 1 - 1 = 0\,. \tag{16}$$

Let's denote

$$\sum_{i\in N^+}(1/N - w_i) = -\sum_{i\in N^-}(1/N - w_i) = \gamma < 0$$
$$\beta = \sum_i(1/N - w_i)a_i < a_{i1}\gamma - a_{i2}\gamma = (a_{i1} - a_{i2})\gamma < 0\,. \tag{17}$$

Therefore, the average allele fitness change after a random SPFL change $\langle\Delta w\rangle_{i,w'} = \beta < 0$.

#### 1.2. The new SPFL is partially correlated with the old one

We shall prove that even if the fitness values after a SPFL change ($w'$) are not completely random, but are correlated with the previous SPFL ($w$), the average change in fitness of the current alleles after a random SPFL change is still negative.

Let's assume that the new SPFL has a random component $w^r$ contributing the fraction $\delta$ ($0 < \delta < 1$) of the SPFL, and the remainder $(1 - \delta)$ comes from the old SPFL:

$$w' = \delta w^r + (1 - \delta)w\,. \tag{18}$$

Equation 7 can then be transformed:

$$\beta = \langle\sum_i(w'_i - w_i)a_i\rangle_{w'} = \sum_i(\langle w'_i\rangle_{w'} - w_i)a_i = \sum_i\langle\delta w^r_i + (1-\delta)w_i\rangle_{w'}a_i - \sum_i w_i a_i =$$
$$\delta\sum_i\langle w^r_i\rangle_{w'}a_i + (1-\delta)\sum_i\langle w_i\rangle_{w'}a_i - \sum_i w_i a_i\,. \tag{19}$$

Since the old SPFL doesn't depend on the new SPFL, and $\langle w_i\rangle_{w'} = w_i$,

$$\beta = \delta(\sum_i\langle w^r_i\rangle_{w'}a_i - \sum_i w_i a_i) = \delta\sum_i(1/N - w_i)a_i\,. \tag{20}$$

Since $\delta > 0$ and, as we already proved, $\sum_i(1/N - w_i)a_i$ is negative (eqn. 17), $\beta$ is also negative.

### 2. SPFL changes at rate comparable with the evolution rate ($\tau_M \approx \tau_L$)

If the SPFL changes are not so rare, the distribution of allele probabilities $a_i$ after a substitution doesn't reach the equilibrium state, meaning that $a_i$ does not necessary increase monotonically with the increase of $w_i$ and that equation 6 is not applicable. However, if there is a correlation between $w_i$ and $a_i$ which is decreased by random changes in SPFL, such changes will still result in a decline in the average fitness of the current allele.

Let us represent the relationship between $w_i$ and $a_i$ as the ratio between the components of $w_i$ and $a_i$ vectors, i. e. covariance $cov(a, w)$. Based on eqn. 1 and eqn. 2,

$$cov(a,w) = \langle w_i a_i \rangle_i - \langle w_i \rangle_i \langle a_i \rangle_i = \\ (1/N)\sum_i w_i a_i - ((1/N)\sum_i w_i)((1/N)\sum_i a_i) = (1/N)\sum_i w_i a_i - (1/N^2)\,. \tag{21}$$

The equation 3 can be transformed:

$$\beta = \langle \Delta w \rangle_{i,w'} = \langle w'_i - w_i \rangle_{i,w'} = \langle \sum_i (w'_i - w_i) a_i \rangle_{w'} = \\ \langle \sum_i w'_i a_i \rangle_{w'} - \langle \sum_i w_i a_i \rangle_{w'} = \langle cov(w', a) \rangle_{w'} - \langle cov(w, a) \rangle_{w'} = \\ \langle cov(w', a) \rangle_{w'} - cov(w, a)\,. \tag{22}$$

If the covariance of the new fitness values $w'_i$ and the probabilities of the alleles to be present $a_i$ is on average lower than it was before the SPFL change, $\beta$ is negative and the current allele fitness on average decreases when the SPFL changes.

We should first prove the existence of a positive correlation between the fitness of an allele and its probability.

If time is viewed as a discrete variable, then the substitution matrix $f$ represents the probability of the allele $j$ to be replaced by the allele $i$ at the next time step:

$$f = \frac{1}{N}\begin{bmatrix} N - \sum_{i\neq 1} f_{i1} & f_{12} & f_{13} & \cdots & f_{1N} \\ f_{21} & N - \sum_{i\neq 2} f_{i2} & f_{23} & \cdots & f_{2N} \\ f_{31} & f_{32} & N - \sum_{i\neq 3} f_{i3} & \cdots & f_{3N} \\ \vdots & \vdots & \vdots & \ddots & \vdots \\ f_{N1} & f_{N2} & f_{N3} & \cdots & N - \sum_{i\neq n} f_{iN} \end{bmatrix}. \tag{23}$$

Here, $f(s_{ij}) = f_{ij}$ is the monotonically increasing function of selection coefficient. We shall prove that under selection, the correlation between fitness vector $w_i$ and the distribution of allele probabilities $a_i$ emerges. For this purpose, we will demonstrate that the frequency distribution at the next time step $a'_i$ will be on average more strongly correlated with $w_i$ than the initial $a_i$:

$$\Delta = \langle cov(w, a') \rangle_a - \langle cov(w, a) \rangle_a \geq 0\,. \tag{24}$$

Based on equations 1 and 2,

$$cov(w, a) = \sum_k w_k a_k - (\sum_k w_k)(\sum_k a_k) = \sum_k w_k a_k - 1\,. \tag{25}$$

We can substitute this expression into the equation 24:

$$\Delta = \langle cov(w, a') \rangle_a - \langle cov(w, a) \rangle_a = \langle \sum_k w_k a'_k \rangle_a - \langle \sum_k w_k a_k \rangle_a\,. \tag{26}$$

The frequency of the allele $k$ at the next time step $a'_k$ is the result of multiplying the matrix $f$ by $a$:

$$a'_k = \sum_j f_{kj} a_j \,. \tag{27}$$

Then

$$\begin{aligned} \Delta = \langle \sum_k w_k a'_k \rangle_a - \langle \sum_k w_k a_k \rangle_a = \\ \langle \sum_k w_k \sum_j f_{kj} a_j \rangle_a - \langle \sum_k w_k a_k \rangle_a = \langle \sum_k w_k \sum_j f_{kj} a_j - \sum_k w_k a_k \rangle_a = \\ \langle \sum_k w_k (\sum_j f_{kj} a_j - a_k) \rangle_a = \langle \sum_k w_k (\sum_j f_{kj} a_j - \sum_j I_{kj} a_j) \rangle_a = \\ \langle \sum_k w_k (\sum_j f_{kj} - \sum_j I_{kj}) a_j) \rangle_a = \langle \sum_k w_k \sum_j (f_{kj} - I_{kj}) a_j \rangle_a = \\ \sum_k w_k \sum_j (f_{kj} - I_{kj}) \langle a_j \rangle_a = \sum_k w_k \sum_j (f_{kj} - I_{kj}))(1/N) , \end{aligned} \tag{28}$$

where $I$ is the identity matrix. Therefore, since $N > 0$, the inequality 24 is transformed to

$$\sum_k w_k \sum_j (f_{kj} - I_{kj}) \geq 0 \,. \tag{29}$$

The matrix in the brackets is as follows:

$$f_{ij} - I_{ij} = \frac{1}{N} \begin{bmatrix} -\sum_{i \neq 1} f_{i1} & f_{12} & f_{13} & \dots & f_{1N} \\ f_{21} & -\sum_{i \neq 2} f_{i2} & f_{13} & \dots & f_{2N} \\ f_{31} & f_{23} & -\sum_{i \neq 3} f_{i3} & \dots & f_{3N} \\ \vdots & \vdots & \vdots & \ddots & \vdots \\ f_{N1} & f_{N2} & f_{N3} & \dots & -\sum_{i \neq n} f_{iN} \end{bmatrix} \tag{30}$$

The inner sum in the equation 29 is the sum of the $k^{th}$ row of this matrix:

$$\begin{aligned} \sum_k w_k \sum_j (f_{kj} - I_{kj}) = \sum_k w_k (-\sum_{i \neq k} f_{ik} + \sum_{j \neq k} f_{kj}) = \\ \sum_k w_k \sum_{i \neq k} (f_{ki} - f_{ik}) = \sum_{ki} w_k g_{ki} \,, \end{aligned} \tag{31}$$

where we denoted the $f_{ki} - f_{ik}$ matrix as $g_{ki}$. The $g_{ki}$ matrix is anti-symmetrical, i. e. $g_{ki} = -g_{ik}$:

$$g_{ki} = f_{ki} - f_{ik} = -(f_{ik} - f_{ki}) = -g_{ik} \,. \tag{32}$$

Therefore, the diagonal values of the $g_{ki}$ matrix are zero.

The equation 31 is the sum of the $g_{ki}$ matrix values with some weights $w_k$. In the trivial case, if the weights are equal, the sum of the anti-symmetrical matrix values will be zero. However, if it's not the case (i. e. the fitness values of the alleles are not equal and the SPFL is not "flat"), the sum will be not equal to zero and, moreover, positive. To prove it, we split the equation 31 by the $g_{ki}$ matrix diagonal:

$$\begin{aligned} \sum_{ki} w_k g_{ki} = \sum_{ki, k<i} w_k g_{ki} + \sum_{ki, k>i} w_k g_{ki} = \sum_{ki, k<i} w_k g_{ki} + \sum_{ik, i>k} w_i g_{ik} = \\ \sum_{ki, k<i} (w_k g_{ki} + w_i g_{ik}) = \sum_{ki, k<i} (w_k g_{ki} - w_i g_{ki}) = \sum_{ki, k<i} g_{ki} (w_k - w_i) \,. \end{aligned} \tag{33}$$

Assume that the fitness vector is sorted highest to lowest:

$$w_1 \geq w_2 \geq w_3 \geq \cdots \geq w_N \,. \tag{34}$$

Then, for each $k$, the selection coefficients $s_{ki}$ and the probabilities of the $k \longrightarrow i$ substitution $f(s_{kj}) = f_{kj}$ are also sorted in a decreasing order:

$$\begin{aligned} s_{k1} \geq s_{k2} \geq s_{k3} \geq \cdots \geq s_{kN}\,, \\ f_{k1} \geq f_{k2} \geq f_{k3} \geq \cdots \geq f_{kN}\,. \end{aligned} \tag{35}$$

The sum in the equation 33 is taken over the upper triangular portion of the $g_{ki}$ matrix and, based on the decreasing order of the alleles fitnesses (equation 35), contains only the positive sums:

$$k < i \Rightarrow w_k \geq w_i \Rightarrow (w_k - w_i) \geq 0 \tag{36}$$

$$k < i \Rightarrow w_k \geq w_i \Rightarrow s_{ki} \geq s_{ik} \Rightarrow f_{ki} \geq f_{ik} \Rightarrow g_{ki} = f_{ki} - f_{ik} \geq 0\,. \tag{37}$$

If there is at least one pair of alleles $i$ and $k$ such that $w_i \neq w_k$, the sum is positive:

$$\sum_{ki,k<i} g_{ki}(w_k - w_i) > 0 \tag{38}$$

From equation 28, $\Delta$ is also positive. In other words, even a single substitution increases the correlation between the fitness of the allele and the probability that it resides at the site; random SPFL changes on average eliminate this correlation, decreasing, on average, the fitness of the current allele ($\beta < 0$).

### 3. Generalization of the frequent SPFL changes case ($\tau_M \approx \tau_L$) accounting for differences in mutation rates $\mu_{ij}$

In the above, we assumed that mutations between different alleles are equiprobable, so that the relative rates of substitutions are only dependent on the corresponding selection coefficients (eqn. 23). If there are differences in the probabilities of $i \longrightarrow j$ mutations ($\mu_{ij}$), the substitution matrix $f$ (eqn. 23) will be transformed:

$$f = \frac{1}{N}\begin{bmatrix} N - \sum_{i\neq 1} f_{i1}\mu_{i1} & f_{12}\mu_{12} & f_{13}\mu_{13} & \cdots & f_{1N}\mu_{1N} \\ f_{21}\mu_{21} & N - \sum_{i\neq 2} f_{i2}\mu_{i2} & f_{13}\mu_{13} & \cdots & f_{2N}\mu_{2N} \\ f_{31}\mu_{31} & f_{32}\mu_{32} & N - \sum_{i\neq 3} f_{i3}\mu_{i3} & \cdots & f_{3N}\mu_{3N} \\ \vdots & \vdots & \vdots & \ddots & \vdots \\ f_{N1}\mu_{N2} & f_{N2}\mu_{N2} & f_{N3}\mu_{N3} & \cdots & N - \sum_{i\neq n} f_{iN}\mu_{iN} \end{bmatrix}. \tag{39}$$

Accordingly, the $g_{ki}$ matrix from eqn. 31 will be equal to $f_{ki}\mu_{ki} - f_{ik}\mu_{ik}$. The $g_{ki}$ matrix will still be anti-symmetrical, i. e. $g_{ki} = -g_{ik}$. However, the $g_{ki}$ will no longer necessarily be positive: it may be negative if the mutation bias and selection act in opposite directions. Indeed, the probability that a site is occupied, at equilibrium, by a deleterious allele can be high if the rate of mutation to this allele is much higher than the rate of mutation from this allele [2, 3].

Nevertheless, we'll prove that the eqn. 38 will still hold when we average across all possible new SPFLs, i. e.

$$\langle \sum_{ki,k<i} g_{ki}(w_k - w_i)\rangle_w > 0, \tag{40}$$

as long as the selection coefficient (corresponding to the new SPFL) and the mutation rate for a pair of alleles are independent of each other:

$$\langle f_{ik}\mu_{ik}\rangle_w = \langle f_{ik}\rangle_w \langle \mu_{ik}\rangle_w = \langle f_{ik}\rangle_w \mu_{ik}\,. \tag{41}$$

Equation 40 can be transformed:

$$
\begin{aligned}
n\langle\Delta\rangle_w &= \langle \sum_{ki,k<i} g_{ki}(w_k - w_i)\rangle_w = \langle \sum_{ki,k<i} (f_{ki}\mu_{ki} - f_{ik}\mu_{ik})(w_k - w_i)\rangle_w = \\
\sum_{ki,k<i} \langle(f_{ki}\mu_{ki} - f_{ik}\mu_{ik})(w_k - w_i)\rangle_w &= \sum_{ki,k<i} \langle f_{ki}\mu_{ki}(w_k - w_i)\rangle_w - \sum_{ki,k<i} \langle f_{ik}\mu_{ik}(w_k - w_i)\rangle_w = \\
&\sum_{ki,k<i} \langle f_{ki}\mu_{ki}(w_k - w_i)\rangle_w + \sum_{ki,k>i} \langle f_{ki}\mu_{ki}(w_k - w_i)\rangle_w = \\
&\sum_{ki} \langle f_{ki}\mu_{ki}(w_k - w_i)\rangle_w = \sum_{ki} \mu_{ki}\langle f_{ki}(w_k - w_i)\rangle_w \,.
\end{aligned} \tag{42}
$$

Since $f_{ki}$ is a monotonic function of $(w_k - w_i)$, they are positively correlated:

$$
\langle f_{ki}(w_k - w_i)\rangle_w - \langle f_{ki}\rangle_w \langle(w_k - w_i)\rangle_w = \langle f_{ki}(w_k - w_i)\rangle_w - 0 = \delta > 0 \,. \tag{43}
$$

Then, based on eqn. 42,

$$
n\langle\Delta\rangle_w = \sum_{ki} \mu_{ki}\delta = \delta \sum_{ki} \mu_{ki} \,. \tag{44}
$$

Since the mutation rates are also positive,

$$
n\langle\Delta\rangle_w = \delta \sum_{ki} \mu_{ki} > 0 \,. \tag{45}
$$

Therefore, we have proven that given any mutation rate matrix, even a single substitution on average (across all possible SPFLs) increases the correlation between the frequency of the allele and its fitness, assuming that selection preferences are independent of mutation biases.

### Supplementary Text 2

### The rate of evolution under fluctuating selection

According to the diffusion fluctuation theory [4–6], if the fluctuations in the SPFL are very rapid, the resulting landscape will be "quasi-neutral". In this case, the substitution rate will be reduced, and not increased, by further increase in the fluctuation rate, ultimately reaching the neutral value. This is not modeled in the Markov chain based approach we used for simulations [7]. We assume that most fluctuation-induced substitutions occur when the SPFL change frequency is lower than the rate of evolution or comparable to it, so our model is still suitable to study evolution under fluctuating selection.
