## Supplementary Figures for "Senescence and entrenchment in evolution of amino acid sites"

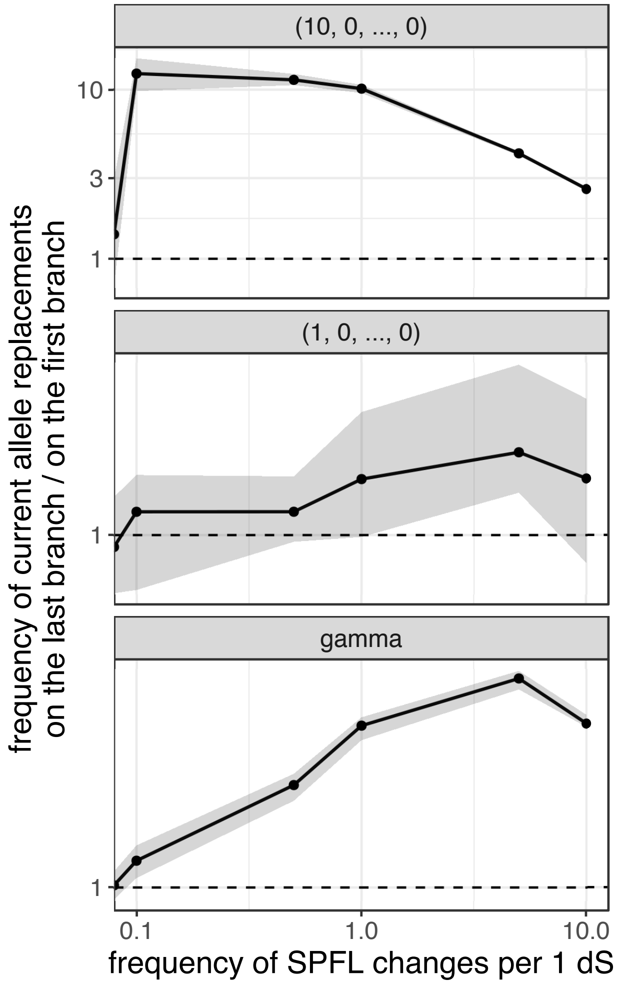


Fig. S1 **Random SPFL changes lead to the increase of the rate of the current allele replacements with the time since its origin**

For the static SPFLs of different shapes, i.e. if initial fitness values are drawn from some distribution and don't change with time, the frequency of the replacements of the current allele remain the same for "early" and "late" branches. If fitness values are redrawn from the same distribution with some frequency (i. e. SPFL changes randomly with time), the rate of current allele replacements on the "late" branch increases. The 95% confidence bands are obtained in 10 simulation repeats.

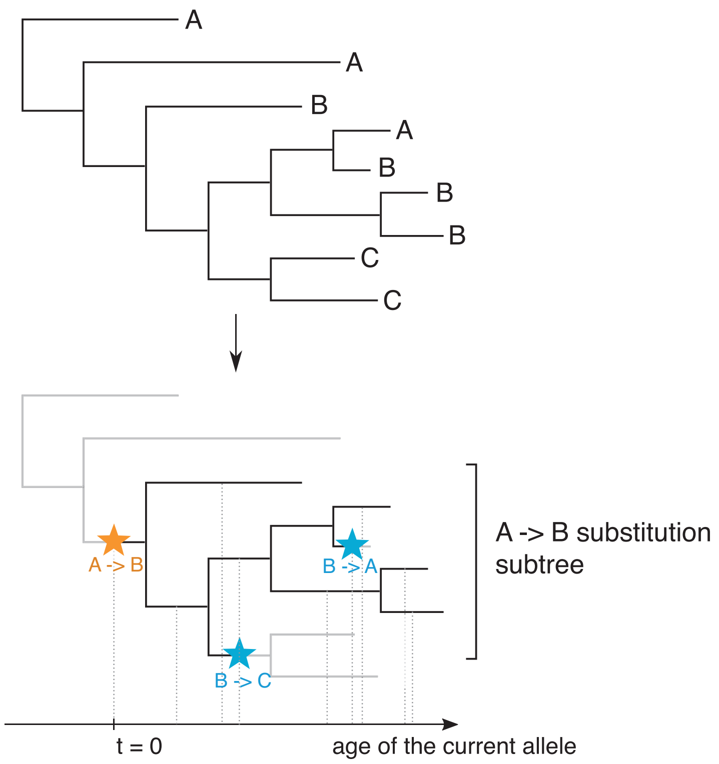


Fig. S2 **The concept of substitution subtrees**
For every individual amino acid site, ancestral state reconstruction can be used to infer allele substitution history at this site. For every replacement $A \rightarrow B$ (orange) on any internal branch of the phylogeny, we can define the corresponding substitution subtree — the contiguous segment of the phylogeny where every internal node carries the derived variant $B$. Within this substitution subtree, the current allele $B$ can be replaced by the ancestral variant $A$ ($B \rightarrow A$ substitution) or some other variant $C$ ($B \rightarrow C$ substitution) in the course of allele losses (blue). For such subtree, $A$ is the ancestral allele and $B$ is the current allele.

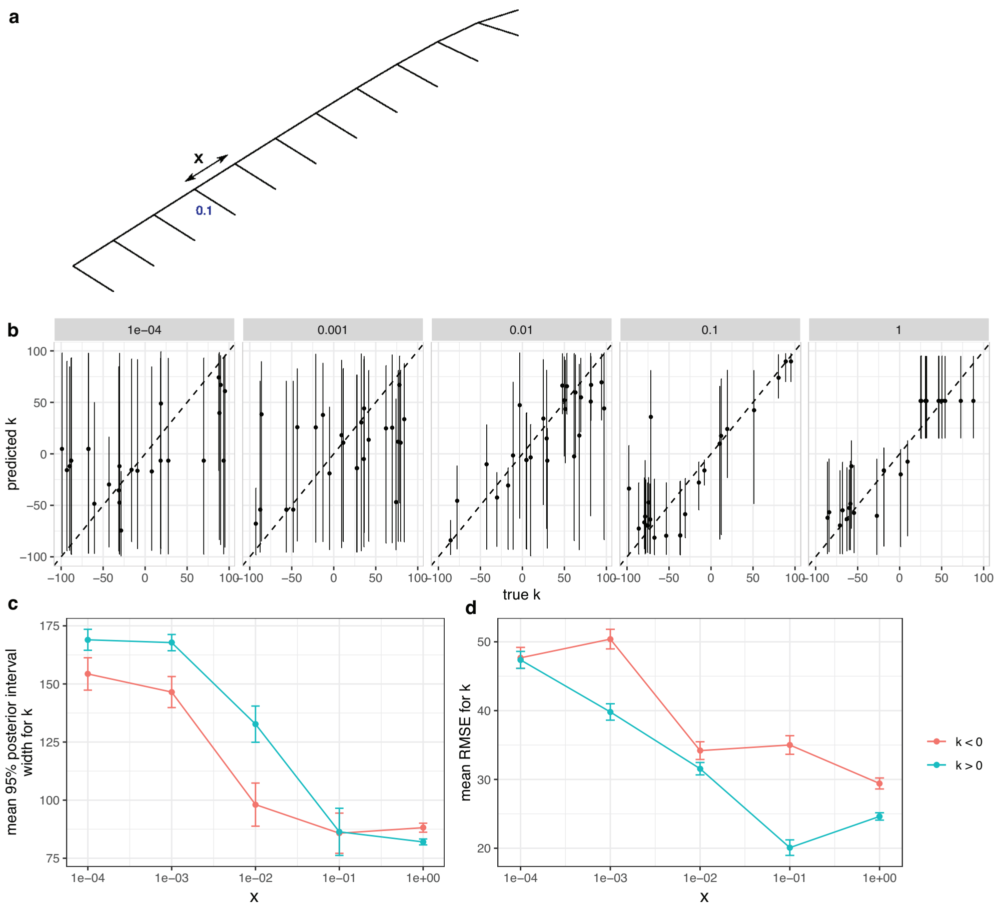


Fig. S3 **Performance of ABC inference of $k$ on phylogenies of different shape**
**a**, The phylogenetic tree used for ABC performance validation. The topology of the phylogeny and the length of the terminal branches (0.1 dS) remained the same, while the length of the internal branches (x) varied from 1e-4 ("star-like" tree) to 1. **b**, The 95% posterior probability intervals for $k$ estimated on the phylogenies with varying $x$; **c**, average width of the 95% posterior probability intervals for $k$ and **d**, average RMSE of the prediction on the phylogenies with varying $x$.

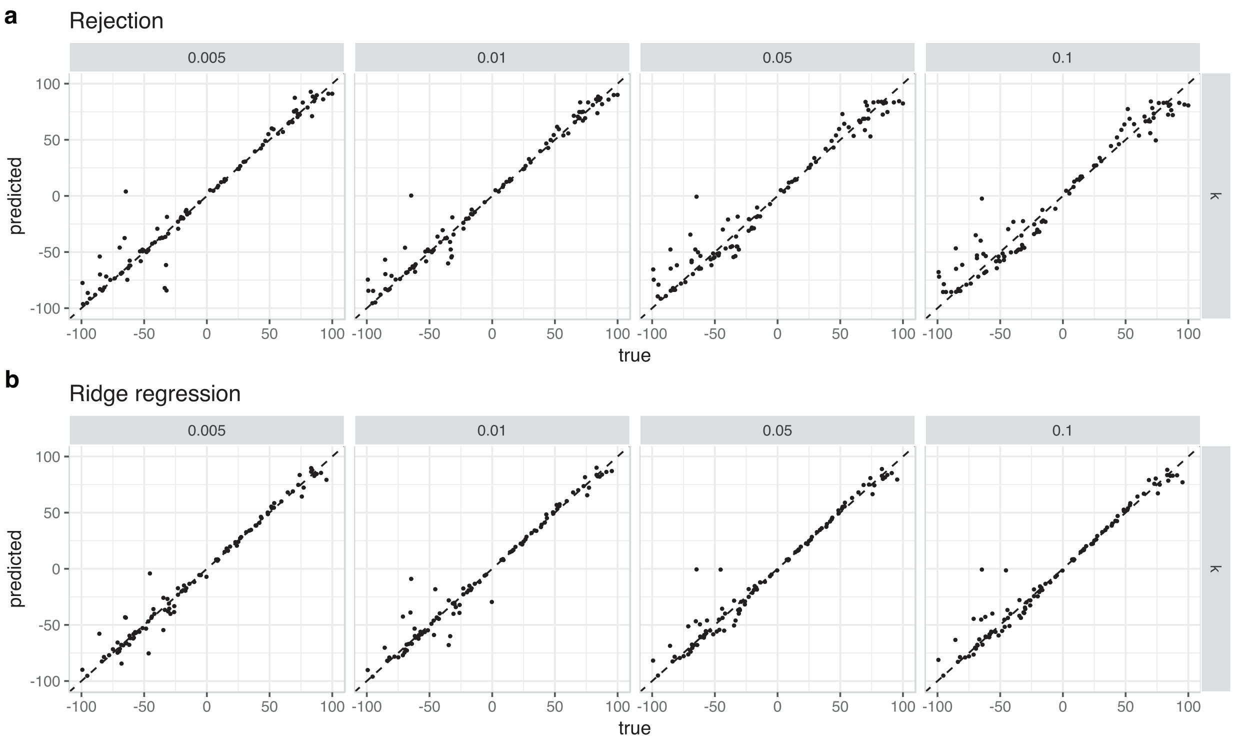


Fig. S4 **Accuracy of the prediction for the two-parameter model**
Estimation accuracy for the rate of the change of the current allele fitness $k$ under 0.005, 0.01, 0.05 and 0.1 acceptance threshold using simple rejection ABC (**a**) and ABC with local Ridge regression adjustment (**b**). The average prediction errors are listed in Table S3.

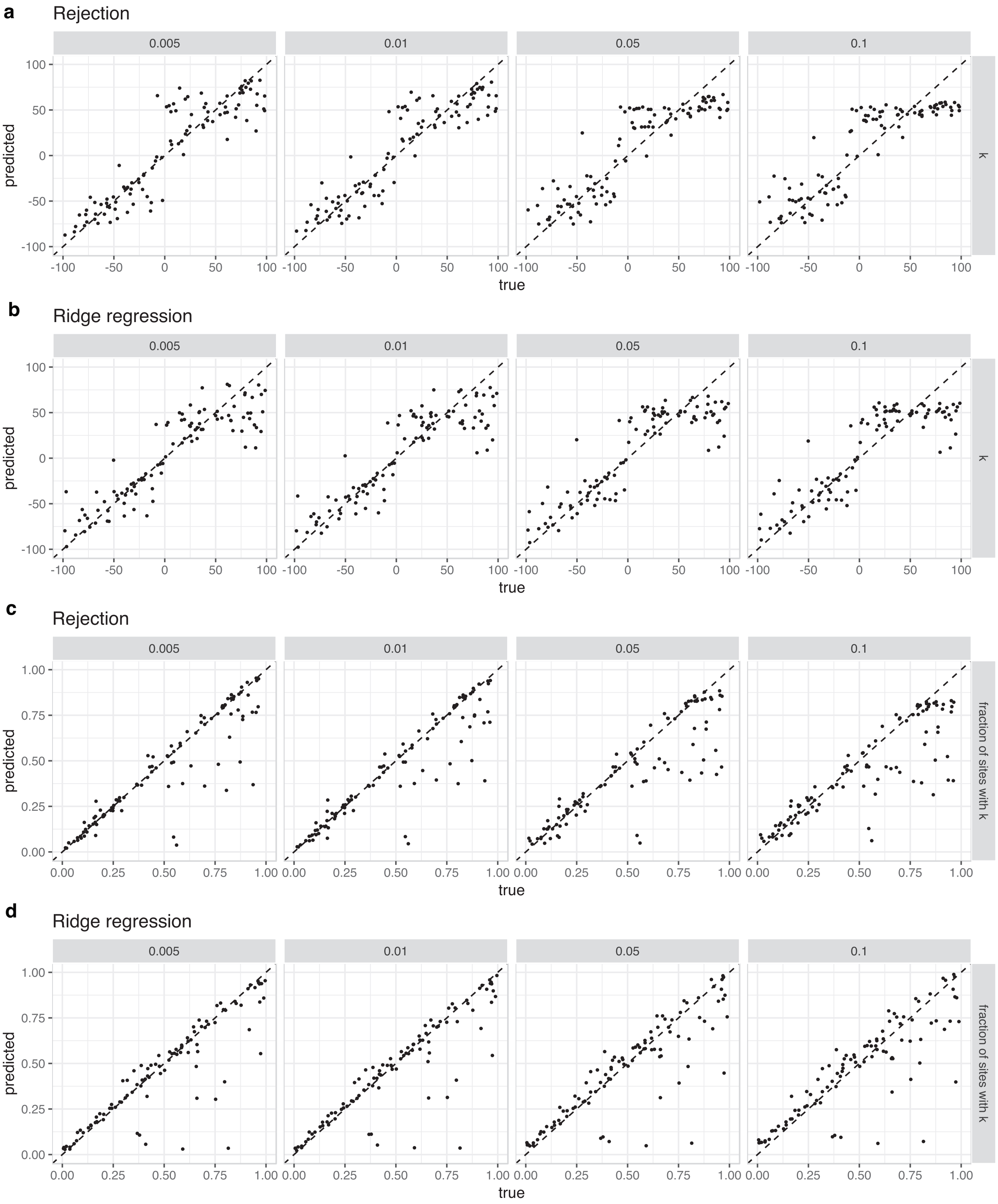


Fig. S5 **Accuracy of the prediction for the three-parameter model**
Estimation accuracy for the rate of the change of the current allele fitness $k$ **(a,b)** and the fraction of alleles with changing fitness **(c,d)** under 0.005, 0.01, 0.05 and 0.1 acceptance threshold using simple rejection ABC **(a,c)** and ABC with local Ridge regression adjustment **(b,d)**. The average prediction errors are listed in Table S3.

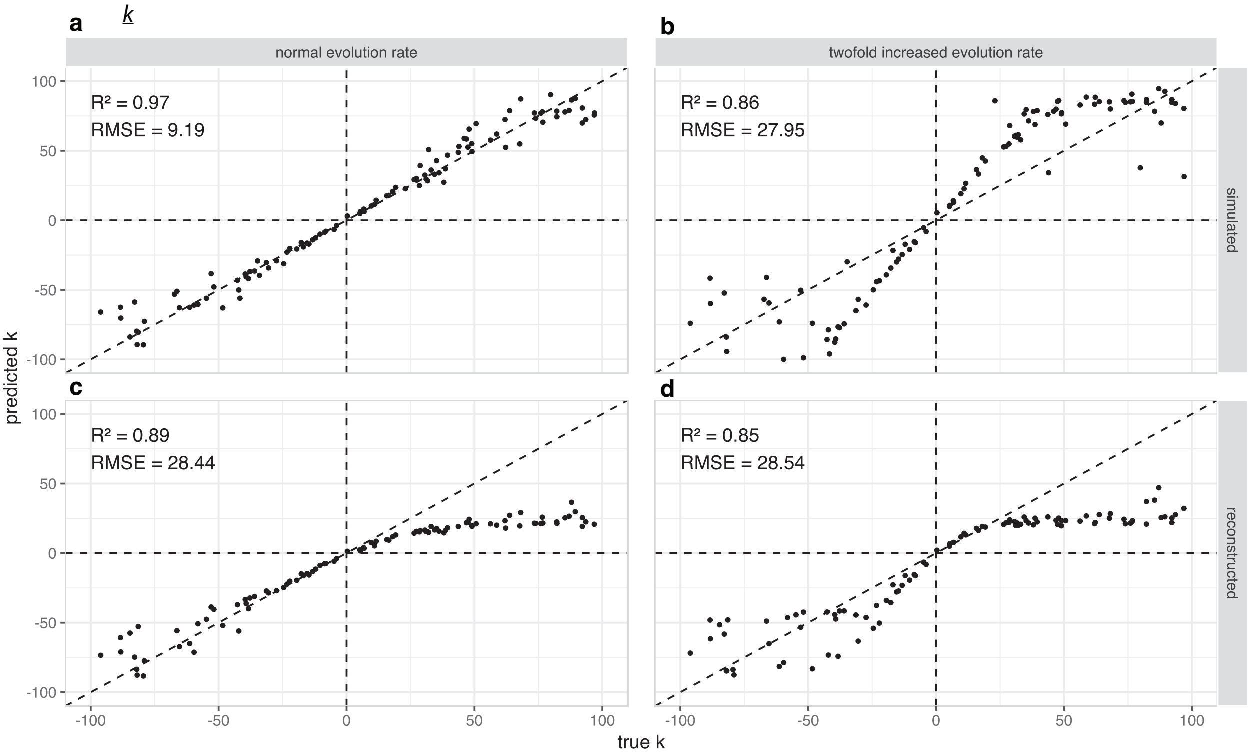


Fig. S6 **Accuracy of the prediction for the two-parameter model under accelerated evolution rate and reconstruction of ancestral states**

The rate of the current allele fitness change $k$ was estimated on simulations with normal (**a,c**) and twofold increased overall substitution rate (**b,d**). For **a,b** the substitution history was taken directly from simulations; for **c,d** it was reconstructed with *codeml*. The confusion matrices for all four cases are shown in Table S2.

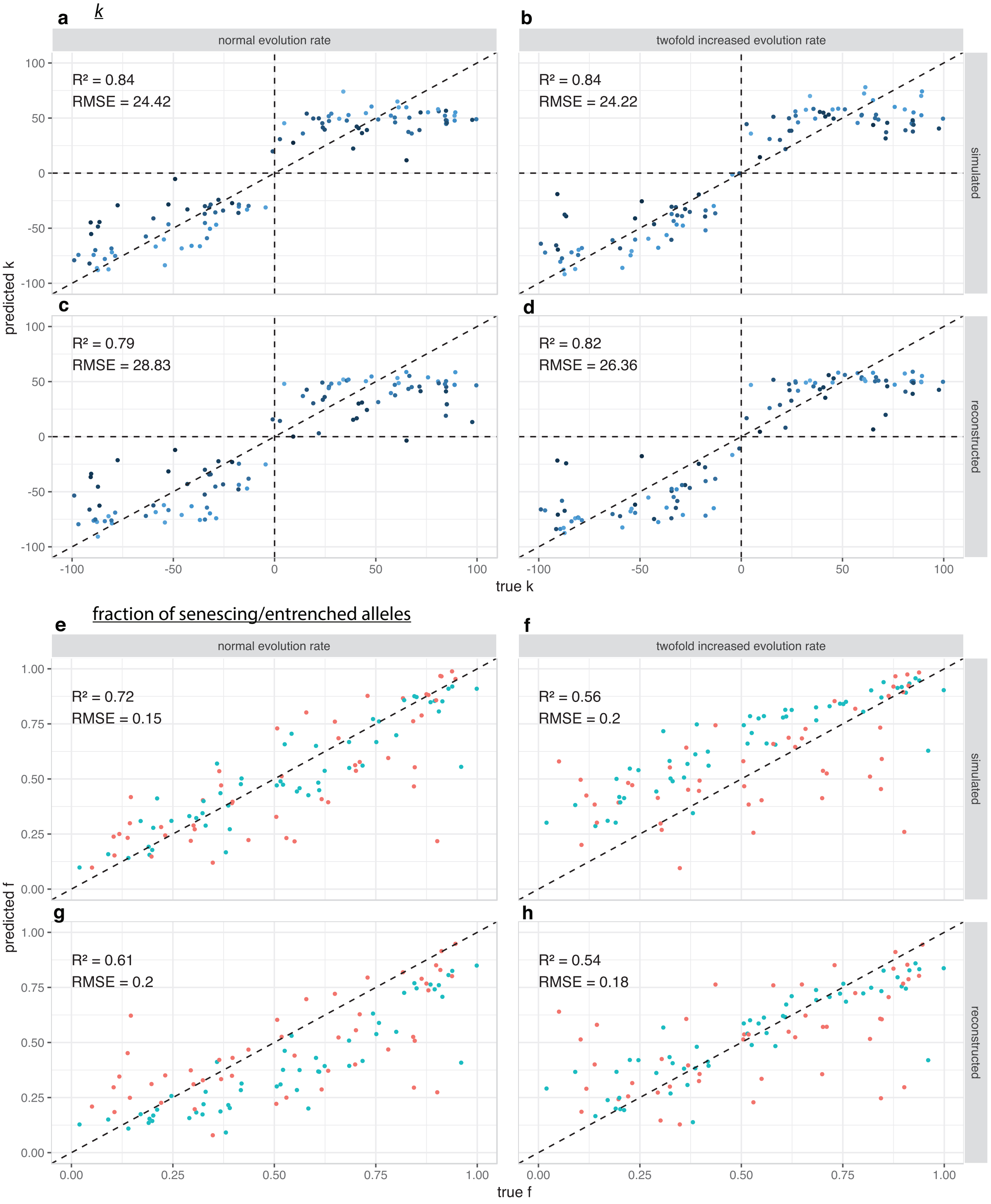


Fig. S7 **Accuracy of ABC prediction for the three-parameter model with and without accelerated evolution rate and reconstruction of ancestral states**

**a-d**, The rate of the current allele fitness change $k$ was estimated on simulations with normal (**a,c**) and twofold increased overall substitution rate (**b,d**). For **a,b** the substitution history was taken directly from simulations; for **c,d** it was reconstructed with *codeml*. The confusion matrices for all four cases are shown in Table S2. **e-h**, The same for the fraction of alleles with changing fitness based on the same set of simulations. Blue dots correspond to the simulations with $k > 0$ (entrenchment), red dots correspond to the simulations with $k < 0$ (senescence).

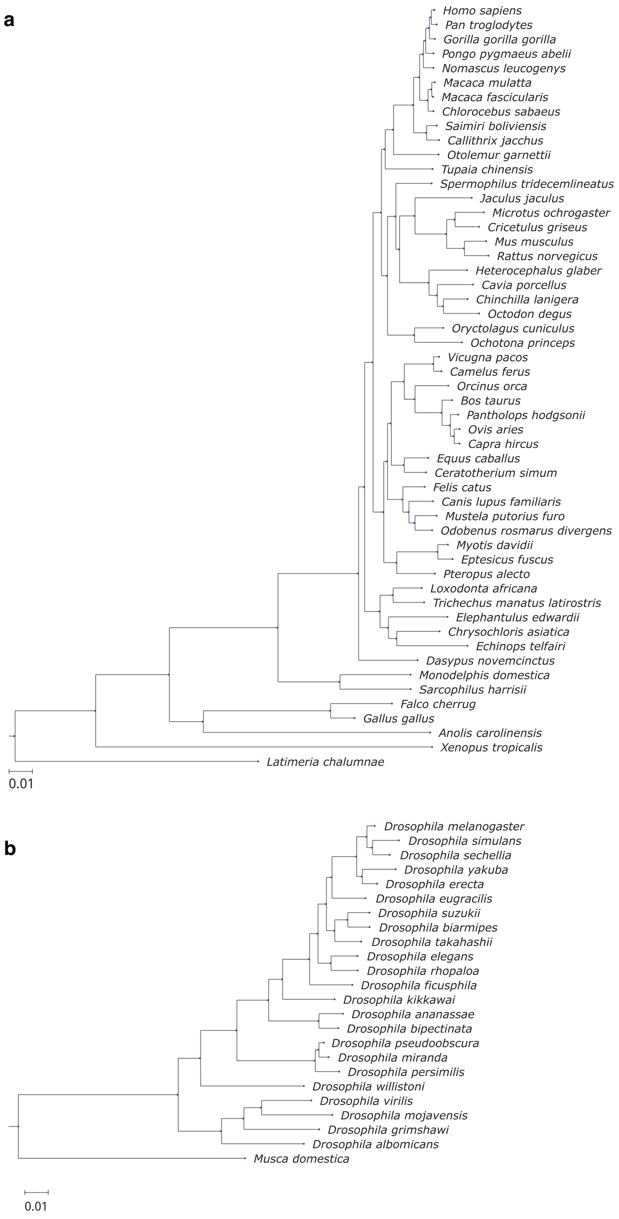


Fig. S8 **The phylogenies used for the inference of current allele fitness change**
The phylogenies of 53 species of vertebrates (**a**) and of 24 species of insects (**b**) from UCSC Genome Browser database. The branch lengths are given in dS.

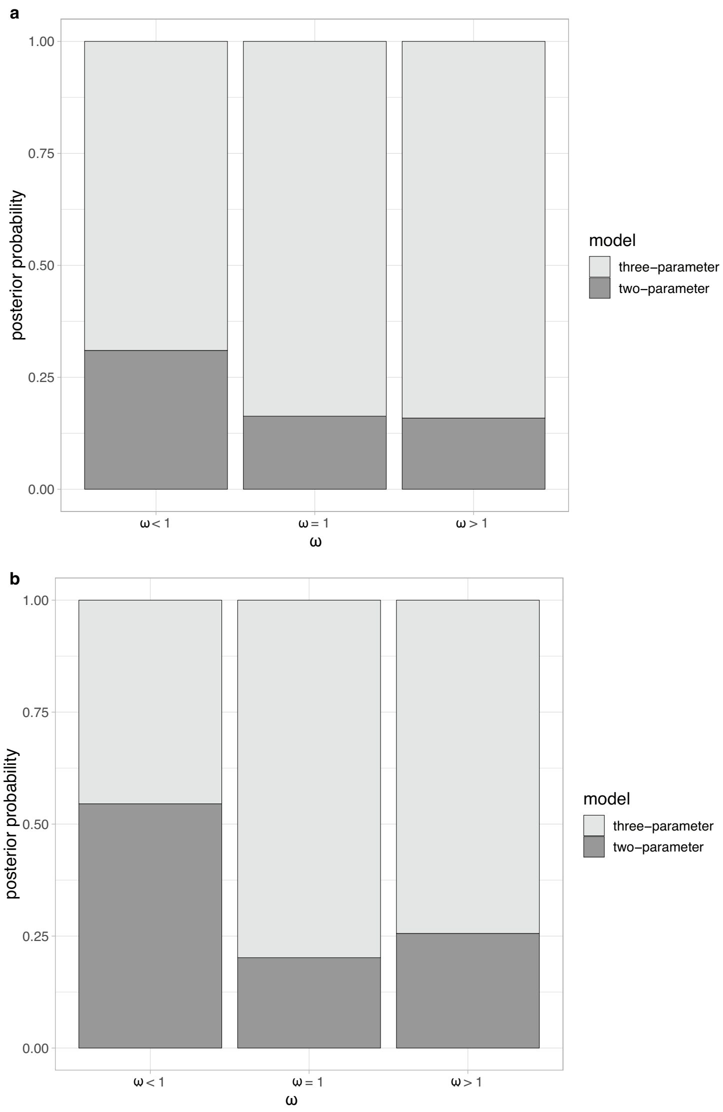


Fig. S9 **The posterior probabilities for the two- and three-parameter models**
The ABC posterior probabilities for the two-parameter (dark grey) and the three-parameter (light grey) models for the vertebrates (**a**) and insects (**b**) data. For all datasets, except sites with $\omega < 1$ of insects, the three-parameter model is preferable.

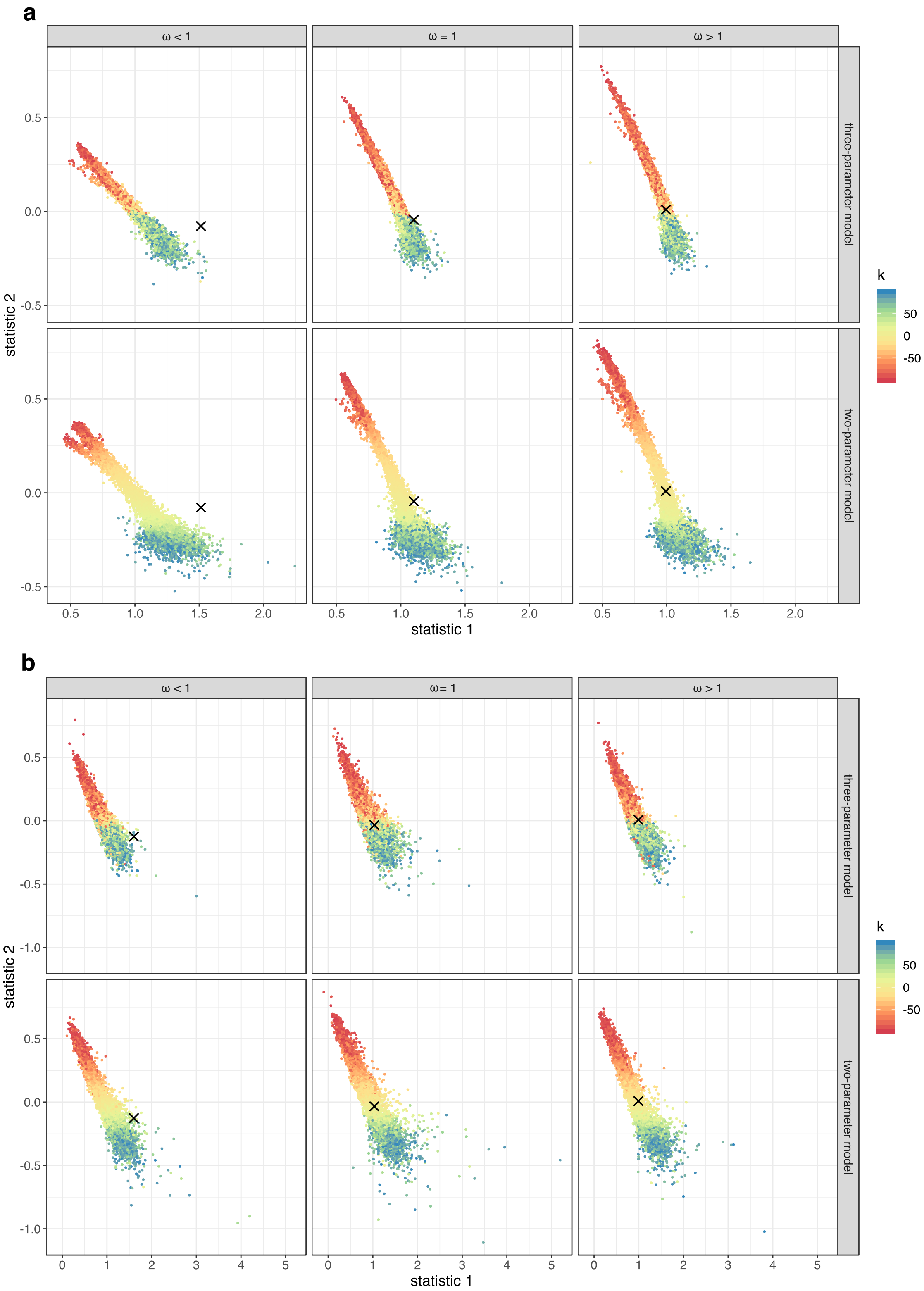


Fig. S10 **The distribution of summary statistics in simulations and data**
Each dot represents one simulation in the space of two summary statistics used in ABC parameter inference. The color shows the value of $k$ used in the simulation (blue corresponds to entrenchment ($k > 0$), red corresponds to senescence ($k < 0$). The distributions for both the two-parameter and the three-parameter models are shown. The crosses show the summary statistics calculated from protein sequences of vertebrates (**a**) and insects (**b**).

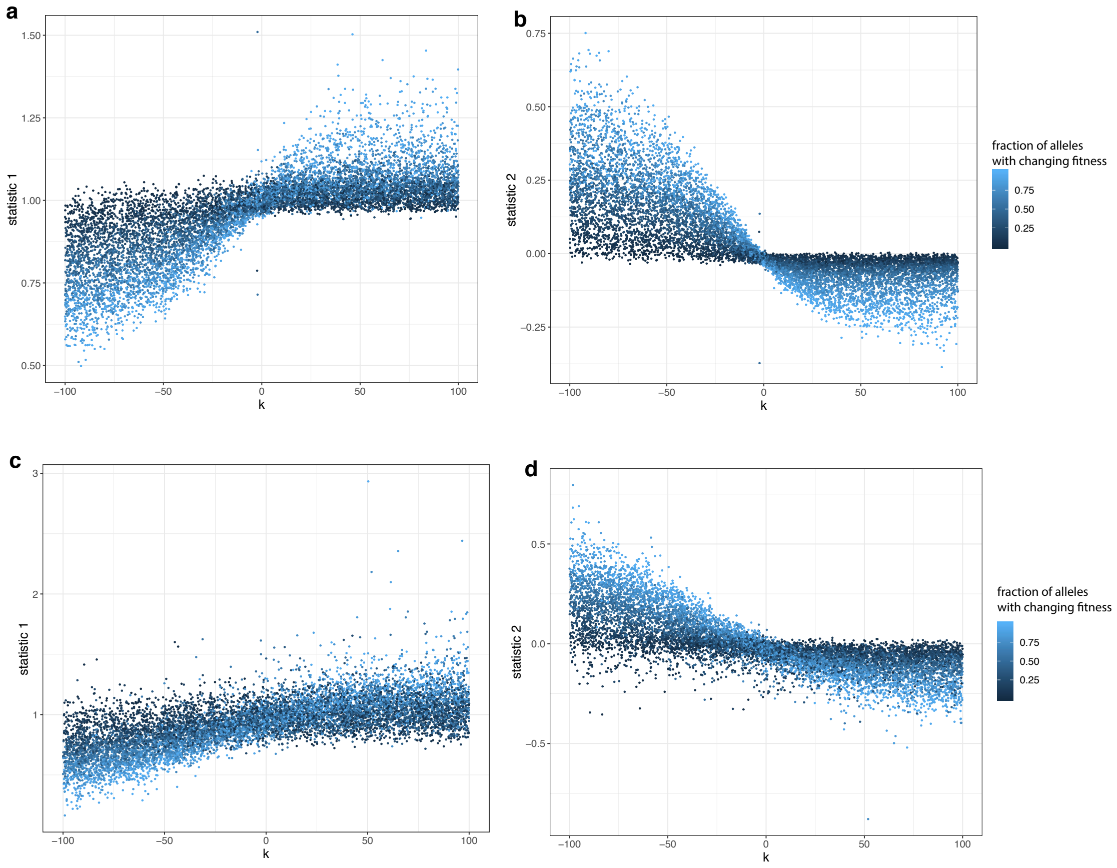


Fig. S11 **The dependence of the summary statistics on the simulation parameters**
The simulations were performed on the phylogenies of vertebrates (**a,b**) and insects (**c,d**). Each dot represents one simulation with some parameters $k$ and fraction of alleles with log fitness linearly changing with the rate $k$ (color). For both phylogenies, summary statistics reflect both the direction and the rate of current allele fitness change $k$ and the fraction of alleles with dynamic fitness.

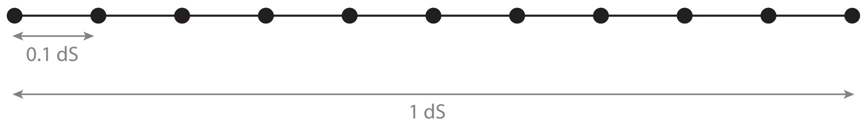


Fig. S12 **The model phylogeny used in simulations**
The phylogeny which was used to demonstrate the effects of senescence, entrenchment and the heterogeneity of the substitution rates.
